# A reference-calibrated spatial-statistical framework for quantifying stroma-tumor cell spatial associations from histology and its geometric limits

**DOI:** 10.64898/2026.10.09.758046

**Authors:** Tatiana Miti, Daria Miroshnycenko, Bina Desai, Andriy Marusyk, David Basanta

**Affiliations:** Department of Integrated Mathematical Oncology, H Lee Moffitt Cancer Center and Research Institute, Tampa, Florida; Department of Metabolism and Physiology, H Lee Moffitt Cancer Center and Research Institute, Tampa, Florida; Laboratory of Systems Pharmacology at Harvard Medical School, Boston, Massachusetts; Department of Molecular Medicine, University of South Florida, Tampa, Florida

## Abstract

Quantifying interactions in biological tissues requires determining whether an interaction exists, the distance over which it operates, and its strength. Existing spatial-statistical approaches can characterize spatial organization but often require interaction range or interaction models to be specified a priori, making them less suited to settings where these quantities are themselves unknown. Here, we present a practical framework that recovers the presence, range, and association strength of spatial bias toward a tissue structure directly from segmented histology. The framework combines four complementary statistics, the resource selection function, nearest-structure distance distributions, the cross J-function, and the radial distribution function, with specimen-specific reference states that preserve tissue stromal architecture while spanning a continuum of known spatial biases. Across controlled *in silico* tissues, we show that tissue stromal architecture imposes geometric limits on the range of recoverable interactions and that event fraction independently constrains statistical sensitivity. These spatial associations establish fundamental bounds on inference and demonstrate that spatial statistics must be interpreted relative to tissue-specific reference states rather than absolute scales. We apply the framework to proliferating tumor cells relative to CAF-rich stroma during tyrosine kinase inhibitor therapy in an ALK+ NSCLC model and identify a strong stroma-associated proliferation bias extending approximately 30-60 µm. The framework is broadly applicable to any histological setting in which events and reference structures can be defined, providing a practical approach for quantifying localized interactions directly from tissue stromal architecture.

## Introduction

Tumors are spatially structured ecosystems in which cancer cells coexist and interact with immune, vascular, and stromal components across multiple interaction ranges^1–5^. As in ecological communities, cellular behavior depends not only on composition but also on spatial context. Whether a cell proliferates, survives therapy, or is eliminated depends on its proximity to signals, nutrients, and physical barriers within the tissue^6–9^. Viewing the tumor microenvironment through this ecological lens motivates analyzing it with the spatial statistics developed to describe how organisms distribute themselves relative to one another and to environmental features. Because these interactions are largely mediated by local signaling and matrix-spatial association processes, they are expected to generate measurable spatial relationships between cellular populations and tissue structures. Characterizing such relationships requires determining whether a spatial interaction *is present*, the *distance* over which it operates, and its *association strength*.

Traditional spatial methods are limited in their ability to recover spatial association range and association strength. Quadrat counts and kernel density estimation require the interaction range to be approximated a priori, and their conclusions depend on that choice^10–12^. Ripley’s K-function aggregates spatial association across distances and does not localize the interaction range^13–15^, whereas Morisita indices quantify clustering without reference to a tissue structure^16,17^ (Supplementary Fig. 2). More recent platforms such as muSpAn^18^ and ATHENA^19^ provide powerful frameworks for multiscale spatial analysis, but they are primarily designed around spatial associations among cell populations and do not explicitly treat continuous tissue structures, such as stroma or vasculature, as reference spatial association partners. Nor do they calibrate inferred spatial associations against the geometric constraints imposed by the observed tissue geometry. Consequently, the quantitative recovery of the presence, range, and association strength of spatial associations relative to those constraints remains largely unexplored.

Another common strategy for analyzing interactions is to fit point patterns extracted from the observed histology to predefined interaction models, including Strauss, Geyer saturation, area-interaction, and soft-core point processes, and infer the parameters governing attraction, inhibition, or clustering^20–23^. While highly effective for testing specific mechanistic hypotheses, these approaches require assumptions regarding the form and scale of the underlying interaction and often rely on conditions such as spatial homogeneity, isotropy, or equilibrium to permit tractable inference. Such assumptions can be difficult to justify in biological tissues, which exhibit pronounced spatial heterogeneity and continuous remodeling during disease progression and treatment. In many biological systems, however, the interaction range and distance-decay profile are themselves unknown and constitute the quantities of interest. Prespecifying an interaction kernel may therefore constrain inference to a limited class of spatial behaviors and obscure alternative interaction structures present in the data.

We therefore developed a practical framework that recovers the presence, range, and association strength of spatial bias toward a tissue structure directly from histology, with the interaction range emerging from the data rather than being assumed. Segmented images are converted into marked spatial point patterns in which one population is treated as the events and a tissue component as the reference structure (Fig. 1d; Fig. 2). For each specimen, we generate two reference states that preserve tissue geometry exactly while permuting only the event labels. The first is a random reference, in which events are not spatially biased. The second is a maximal-bias reference, in which events are placed as close to the structure as the tissue geometry allows. Together, these define the achievable limits of the signal imposed by the observed tissue geometry. Exponential distance gradients of known interaction range provide calibration patterns spanning a continuum of spatial biases between these two reference states (Fig. 3). These gradients are not intended as biological interaction models. Instead, they serve as reference patterns with known spatial scales against which we can evaluate statistical behavior. We interpret four complementary statistics, the resource selection function, nearest-structure distance distributions, the cross J-function, and the radial distribution function, relative to these references to infer whether a spatial interaction is present, over what distance it acts, and how strongly. Because such inference is not guaranteed to succeed, we systematically map how each statistic varies with tissue geometry and event fraction, delineating the regimes in which it can and cannot be trusted. This framework calibrates inferred spatial relationships against the geometric constraints imposed by the observed tissue geometry, allowing interaction association strength and range to be interpreted relative to the tissue’s physical limits.

**Figure 1.**
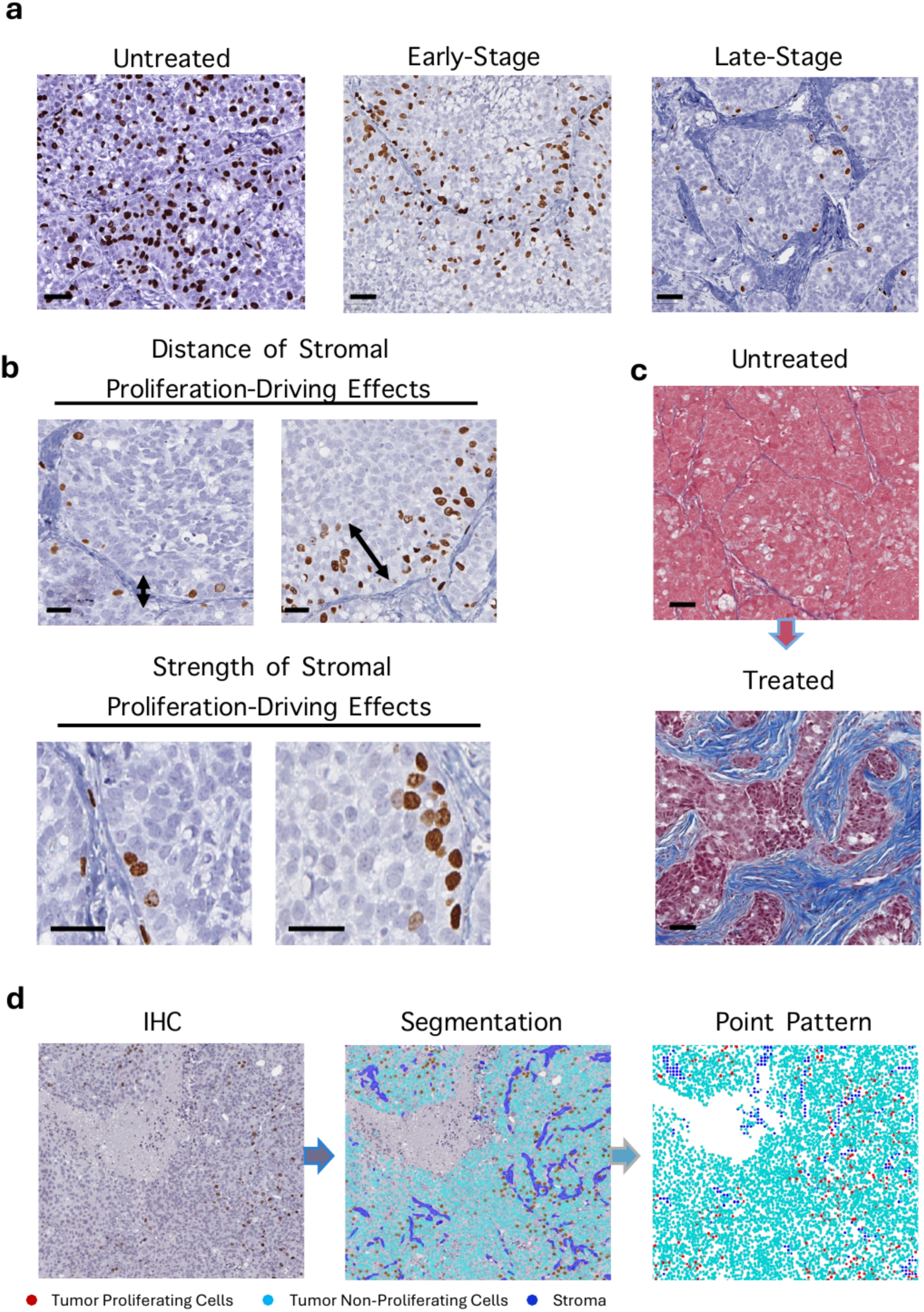
Stroma preserves tumor-cell proliferation during treatment. (a) Representative BrdU immunohistochemistry of ALK+ NSCLC tumors in untreated, early-stage treatment, and late-stage treatment conditions. BrdU+ (proliferating) tumor cells are brown. (b) Higher-magnification fields illustrating the distance over which stromal proliferation-driving effects extend (top) and the strength of the stromal-associated proliferative enrichment (bottom). (c) Masson’s trichrome staining of untreated (top) and treated (bottom) tumors, illustrating treatment-associated stromal remodeling; collagen-rich stroma is blue. (d) Analysis workflow from IHC image to digital segmentation and marked point pattern, with proliferating tumor cells (red), non-proliferating tumor cells (cyan), and stroma (blue). Scale bars are 50 µm in panels a, b, and d; scale bars in panel c are 200 µm.

**Figure 2.**
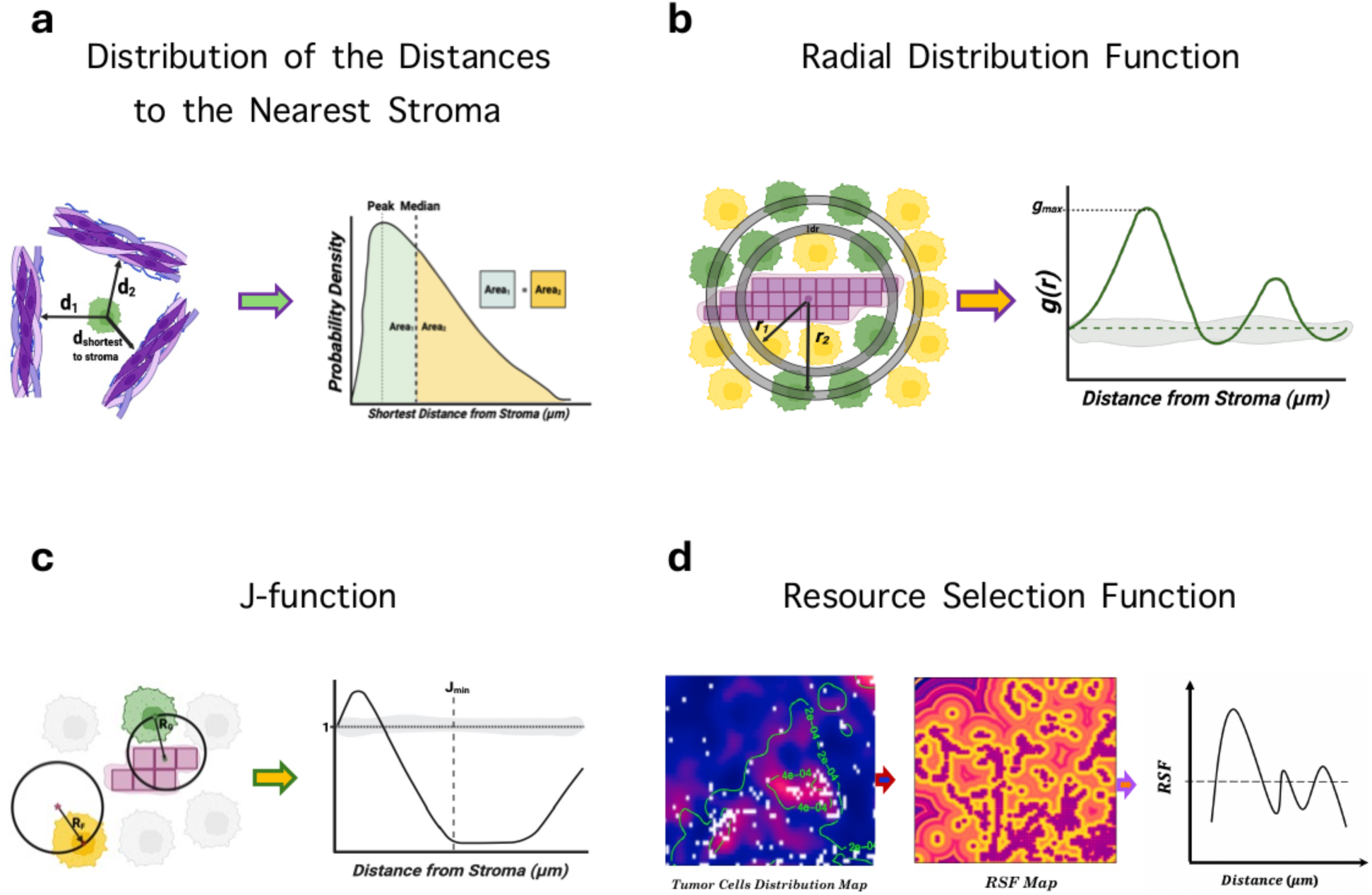
Spatial statistical methods adapted to quantify stroma-tumor-cell spatial associations. (a) Nearest-stroma distance distribution where the shortest distance from each tumor cell to stroma is measured to generate a distance distribution; the schematic illustrates the peak, median, and distributional areas, while quantitative analyses in this study use the median and the KS statistic relative to the random reference. (b) Radial distribution function (RDF), g(r), quantifying enrichment or depletion of proliferating tumor cells at distance r from stroma; maximum RDF (gmax) summarizes the strongest local enrichment. (c) Cross J-function, for which values below 1 indicate greater proximity to stroma than expected under the random reference; Jmin denotes the J value at 60 µm used as the scalar summary in this study. (d) Resource selection function (RSF), showing the tumor-cell distribution map, the distance-to-stroma covariate/RSF map, and the resulting distance-dependent RSF curve.

**Figure 3.**
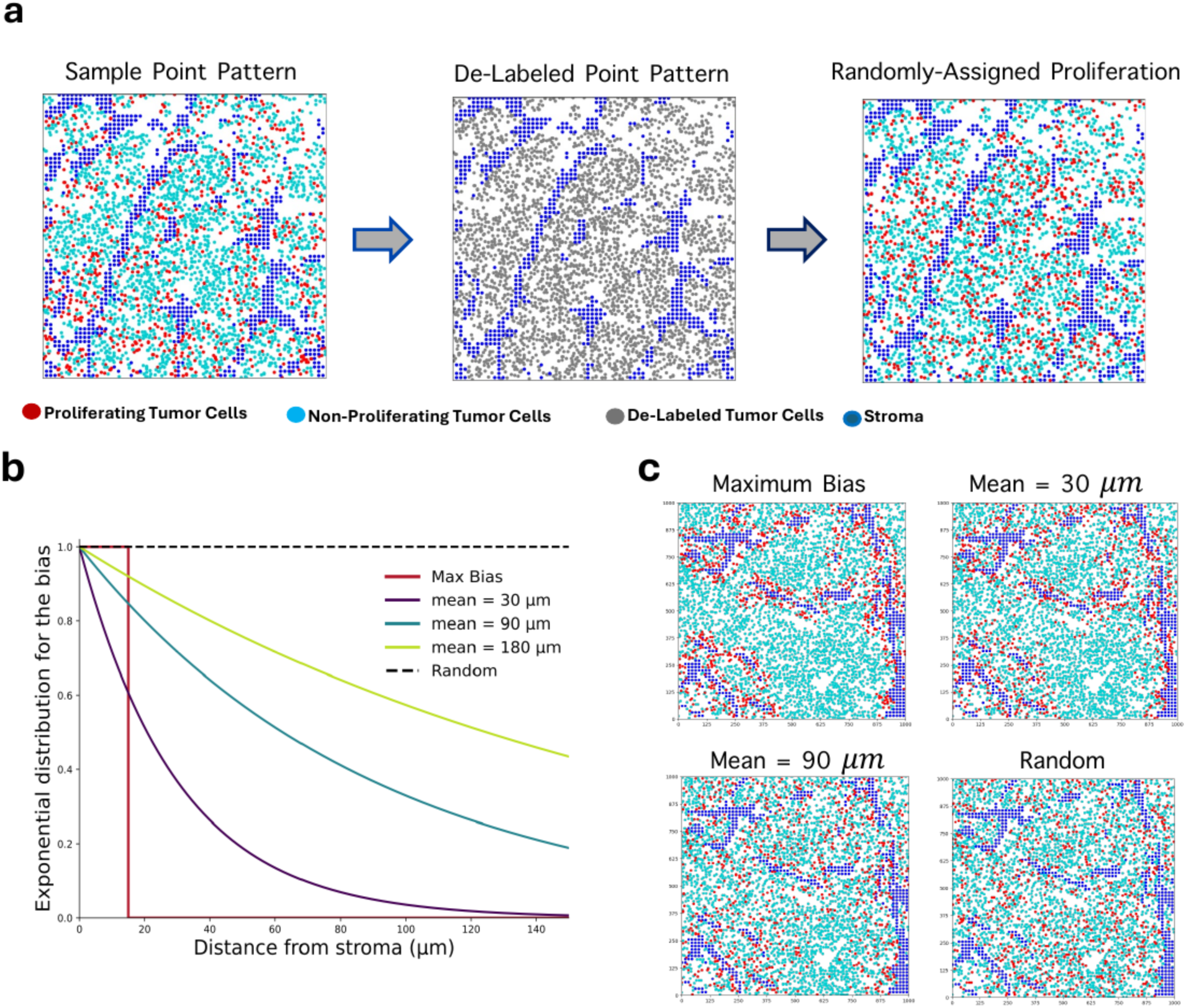
Construction of in silico reference controls for in vivo samples. (a) Reference-state construction where the sample point pattern is de-labeled while retaining all tumor-cell and stromal positions, after which proliferative labels are reassigned according to the specified reference rule. (b) Distance-dependent assignment weights used to generate the reference ladder consisting of a maximal bias, exponential gradients with characteristic distances of 30, 90, and 180 µm, and random/no bias. The maximal-bias curve is shown schematically as the limiting near-stromal assignment; in the simulations, maximal bias is implemented by assigning proliferative labels to the closest available tumor-cell positions until the required proliferative fraction is reached. (c) Representative realizations of maximal bias, mean 30 µm, mean 90 µm, and random reference states on the same stromal and tumor-cell geometry. Proliferating tumor cells are red, non-proliferating tumor cells cyan, de-labeled tumor cells grey, and stroma blue.

We demonstrate the framework using proliferating tumor cells and CAF-rich stromal regions during targeted therapy, a biologically important setting in which localized stromal support has been proposed to contribute to treatment resistance^24–27^. Cancer-associated fibroblasts can adopt therapy-induced pro-tumorigenic states that promote tumor persistence through metabolic support, secretion of survival signals, extracellular matrix remodeling, and other microenvironmental mechanisms. Such spatial associations are expected to act locally through paracrine and matrix-mediated interactions and should therefore generate measurable spatial biases between proliferating tumor cells and stromal tissue. These interactions are particularly relevant in oncogene-driven lung cancers treated with targeted therapies. Although the ALK inhibitor Alectinib induces strong initial responses in ALK-rearranged NSCLC, relapse is nearly universal^28–30^. Using an ALK+ NSCLC mouse model, we examine whether stromal tissue provides a localized treatment-sheltering spatial association that sustains tumor cell proliferation during tyrosine kinase inhibitor therapy and whether this spatial association can be quantified from histology. This setting provides a stringent test of the framework because stromal architecture remodels substantially during treatment, while the proliferative fraction collapses from approximately 50-60% in untreated tumors to approximately 20% within 48 hours and often to 1% or less at treatment nadir. Any quantitative approach must therefore remain interpretable across substantial changes in both stromal organization and event fraction.

The framework applies broadly to any histological setting in which events and reference structures can be defined, including tumor-infiltrating T cells relative to tumor nests, hypoxic cells relative to vessels, and many other examples. This work is deliberately a practical implementation guide rather than a theoretical treatment. We do not provide formal proofs of estimator properties or a full eco-evolutionary model of stromal sheltering. Instead, we focus on constructing the reference states, applying the four statistics, and interpreting them within the geometric limits imposed by tissue geometry and event fraction.

## Results

### A reference-calibrated framework turns descriptive spatial statistics into interpretable measurements of stroma-driven proliferation

The four statistics at the core of this framework, the resource selection function (RSF), nearest-stroma distance distributions, the cross J-function, and the radial distribution function (RDF), are all established estimators available in spatial statistics. None of them, applied on its own, answers the biological question, because each returns a number whose meaning depends on the tissue it was computed in; the same value can reflect a strong spatial association in one specimen and a weak spatial association in another. Our contribution is not a new estimator but a calibration that makes these estimators interpretable, providing a pair of biologically grounded reference states that bound what any statistic can express in each specimen, together with a graded synthetic ground truth against which each statistic’s sensitivity can be read (Figs. 1d, 2, 3).

Histological sections were segmented into proliferating^31,32^ (BrdU+) tumor cells, non-proliferating (BrdU-) tumor cells, and stroma, and converted into a marked point pattern in which tumor cells are nuclear centroids and stroma, a continuous scaffold rather than discrete cells, is discretized onto a ∼15 µm grid (Fig. 1d; Supplementary Fig. 1). The design choice that anchors everything downstream is that we permute proliferative *labels*, never cell *positions*, hence tissue stromal architecture, stromal architecture, and proliferative fraction are held exactly as observed, so any signal a statistic reports is attributable to the spatial arrangement of proliferation alone (kept constant at 20% for all conditions) and not to the fixed stromal architecture.

On this substrate, we build two reference states that bracket the achievable signal in each specimen, and between them a graded ground truth (Fig. 3). The **random reference** redistributes proliferative status at random, defining random reference, the value each statistic takes when there is no stromal bias, and hence the zero point against which *presence* is judged. The **maximal-bias reference** assigns proliferative status to the tumor cells closest to the stroma; it is best understood not as a distance but as the stroma’s *carrying capacity for proliferation*, the maximum proliferation the specimen’s stromal architecture can physically support near it. Within these bounds, we simulate **exponential stromal-spatial association gradients** with a known interaction range of 30-320 µm (Fig. 3b,c), motivated by paracrine factors that diffuse from the stroma and are consumed and degraded. Having all three (a null, a maximal-bias reference ceiling, and a ladder of known-range gradients) and applying them to *every* statistic is the framework’s central advantage. The null gives each statistic a specimen-specific zero, the maximal-bias reference a specimen-specific maximum, while the gradients act as rulers of known length that reveal what a statistic *should* read at a given range. Together, they turn each descriptive curve into a calibrated measurement whose sensitivity, resolution, and failure modes can be read off directly rather than assumed, and they make the four statistics commensurable, so that their agreement becomes meaningful.

Because maximal-bias reference measures an amount rather than a distance, its interaction range depends on stromal content, so only the reference states are legible. Where stroma is scarce, filling the carrying capacity forces proliferating cells outward, so a short-range exponential gradient, one whose interaction range places most of its cells within ∼30-45 µm of stroma, can be more spatially localized than maximal-bias reference itself. On distance-based summaries such as the median and peak location, the expected ordering between the two then reverses, with the short-range gradient sitting closer to stroma than the maximal-bias reference; read without the maximal-bias reference in hand, that smaller median would be mistaken for a stronger spatial association rather than recognized as a broad carrying capacity. This is precisely why every statistic is interpreted by its position within the specimen-specific random-to-maximal-bias reference interval rather than on an absolute scale.

The **nearest-stroma distance distribution** is the histogram of distances from each proliferating cell to its nearest stroma, read against the same distribution for the random reference. When proliferation is stroma-driven, the distribution of proliferating cells shifts toward shorter distances and piles up against the stromal boundary; as spatial association weakens or the proliferative fraction rises, it broadens and shifts rightward until, under the random reference, it matches the control. Two scalars summarize this shift. The **median** is the distance below which half the proliferating cells lie, a robust “typical distance to stroma” that decreases as spatial association strengthens and is unaffected by a handful of far-flung cells. The **KS statistic** is the largest vertical gap between the proliferating and random cumulative curves^33^; it measures the maximum fraction of proliferating cells displaced toward the stroma relative to chance, near 0 when the two distributions coincide, approaching ∼1 when almost every proliferating cell sits closer to the stroma than a random cell would. Because KS integrates the entire distributional shift rather than a single moment, it is the most sensitive test for the *presence* of stroma proliferation-driving spatial associations, whereas the median is the most reproducible. The KS statistic is calculated using two reference populations. In the main figures, we compare BrdU+ cells with the random-label reference, whereas in the supplementary distance-summary figures, we compare BrdU+ cells with non-proliferating (BrdU-) tumor cells.

The **resource selection function (RSF)** plots proliferative intensity against distance to stroma, expressed relative to what spatial availability alone would give, so the curve reads like a profile of spatial association radiating outward from the stromal edge^34,35^. A tall, narrow peak hugging the boundary and returning to baseline (which crucially, is the random reference, not the tissue average) within a few tens of micrometres signals a strong, short-range spatial association; a lower, broader shoulder that only relaxes to baseline at 100-200 µm signals a weaker but longer reaching one; a flat line at baseline means proliferation is placed no more favorably than chance. Two features summarize the curve, specifically the distance at which it returns to baseline estimates the *range*, and its height above baseline estimates the *association strength*. To compare the RSF across conditions as a single scalar, we summarized each curve by its integrated absolute departure from baseline, specifically, the area between the measured relative-intensity curve and the flat no-selection baseline (Methods). Because the baseline is the intensity expected if proliferation ignores stromal distance, this index measures, in one number, how much proliferative intensity departs from that flat expectation across all distances, an association strength readout of the total distance-spatial association selection rather than of any single point on the curve. Unlike the RSF’s peak height or peak location, which are sensitive to the smoothing bandwidth and vary between replicates, the integral averages over the whole curve and is stable, complementing the KS statistic’s test of significance with a measure of association strength.

The **cross J-function** compares how close proliferating cells actually sit to the stroma with how close they could sit given the stroma available, which automatically discounts the fact that stroma-rich tissue makes everything look close^36^. It equals 1 when proximity is no better than chance and falls below 1 when proliferating cells are genuinely nearer to stroma than availability predicts; the distance at which it reaches ∼0 is the range within which essentially every proliferating cell is captured by stroma. A curve that plunges to zero within 40-50 µm means a tight, short-range spatial association; one that eases down over a longer span, or barely departs from 1, means a broad or negligible one. Its scalar is **Jmin** (the J-value at 60 µm) and reads directly as the share of stromal “reach” left unrealized at that 60 *μm* from stroma.

The **radial distribution function (RDF)**, *g*(*r*), is the enrichment of proliferating cells at each separation *r* from stroma expressed as a fold-change over random^23^ where *g(r)* = 1 is chance, *g(r)* > 1 is enrichment, *g(r)* < 1 is depletion, so *g*(45 µm) = 2.7 means 2.7-fold more proliferating cells sit 45 µm from stroma than expected. Its shape distinguishes mechanisms that the cumulative statistics blur together. A curve that is highest right at the boundary and decays monotonically indicates cells decorating the stroma, with enrichment strongest at the shortest distances; the maximal-bias reference takes this form, plateauing near the boundary over the range of the stroma’s *carrying capacity for proliferation* rather than peaking. Its scalar, ***g*max**, is the strongest enrichment reached anywhere on the curve, the cleanest single number for *association strength*; but because it is a density ratio estimated from few points, it becomes noisy when proliferating cells are sparse and must always be read against the random reference.

Because these methods fail under different conditions, the RDF when proliferation is sparse, the cumulative statistics when the tissue geometry truncates long-range separations, the median when the distribution is multimodal, and so on, their agreement guards against artifacts of any one statistic, and mapping *where* each fails, as a function of tissue geometry and proliferative fraction, is itself a central result.

### Pointwise and global envelopes

Both envelopes compare an observed curve against the same null of random labeling, in which BrdU+ status is permuted among the tumor cells while their positions and the stroma are held fixed 99 permutations); the two differ in how they summarize that comparison. The pointwise envelope takes the 2.5th and 97.5th percentiles of the simulations separately at each distance, so the band shows, distance by distance, where the observed curve rises above or falls below what random labeling produces. This is what makes it useful for reading off the range over which stroma affects proliferation, and it is the version we display in the main figures. It is exact, however, only for a test at a single distance chosen in advance, meaning when the curve is compared with the band at every distance at once, under the null it crosses the band somewhere with high probability, which inflates the false-positive rate for any claim about the curve as a whole. The global envelope is built for that whole-curve claim. By ranking the simulated curves rather than treating each distance on its own (the rank-envelope test, GET^37^it yields a single band with simultaneous coverage over the entire tested range and one global p-value, so a curve that leaves the band at any distance is significant at the stated level. We use the two together, this way the pointwise envelope in the main figures to show where proliferation departs from random, and the global envelope for the whole-curve p-values reported throughout. Because the global test is computed on the cumulative distance distribution F(r), the nearest-stroma distance appears as F(r) in the global panels.

### In low-content and dispersed stromal architecture, all four statistics recover both the range and the association strength of stromal proliferation-driving spatial associations

We first present a specimen with low stromal content but high dispersity (247 stromal pixels arranged as thin, dispersed ridges; stroma edge fraction 0.92; Fig. 4f-j; Supplementary Fig. 4). Because the stromal structures are widely spaced, the measurable stromal-distance range extends beyond 200 µm, far larger than in stroma-dense tissue, making this a near-ideal operating regime in which all four methods resolve short-, intermediate-, and long-range gradients.

**Figure 4.**
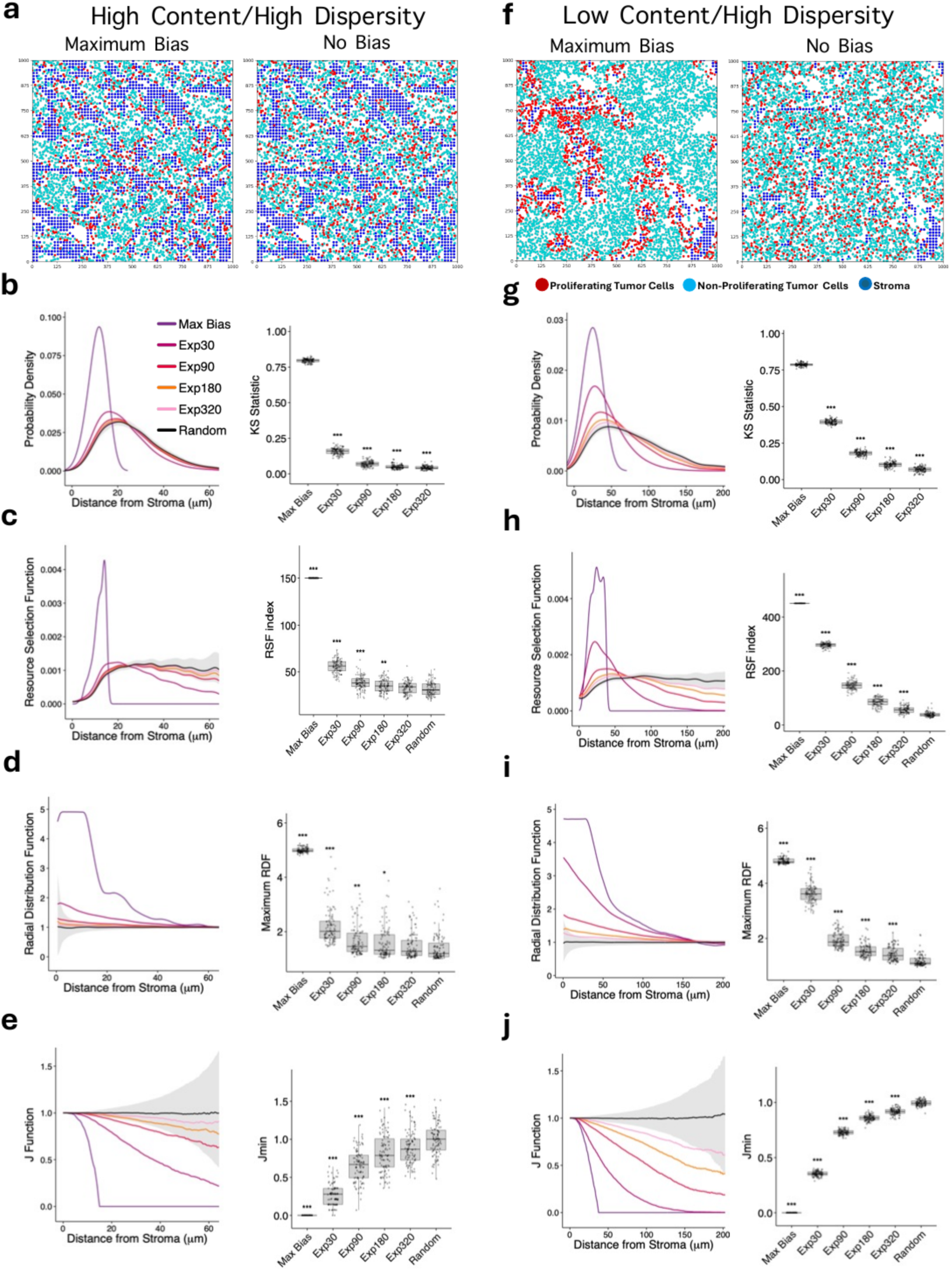
Stromal topology sets the accuracy and sensitivity of spatial analysis. High-content/high-dispersity (a-e) and low-content/high-dispersity (f-j) tissues are analyzed across the same reference-state ladder. (a,f) Representative point patterns for maximum bias and no bias/random reference. (b,g) Nearest-stroma distance probability density (left) and KS statistic (right). (c,h) Resource selection function (left) and RSF index (right). (d,i) Radial distribution function (left) and maximum RDF (right). (e,j) Cross J-function (left) and Jmin (right). Curves compare maximum bias, Exp30, Exp90, Exp180, Exp320, and random/no bias; grey bands show the random-reference envelope where displayed. Points/boxplots show replicate distributions, and significance annotations are shown relative to the random reference as plotted.

Across conditions, the measured RSF curves exhibited the diffusion-like ordering expected for this dispersed, low-content stromal architecture (Fig. 4h). Even when a longer-range spatial association placed its proliferative enrichment farther from the stroma and over a wider band, the stromal architecture yielded good resolution where each curve rose to a near-boundary maximum and decayed monotonically to the flat random baseline, with the decay reaching farther from the stroma as the gradient’s interaction range increased. The maximal-bias reference was the most sharply localized, its enrichment decaying by ∼45-50 µm; the 30 µm gradient peaked near 22 µm and returned to baseline by ∼65-70 µm; and the 90 µm gradient gave a broader but still clearly resolved peak (baseline ∼80 µm). Critically, even the 180 and 320 µm gradients retained weak but positive deviations above a stable random baseline, the long-range discrimination that wide inter-stromal spacing enables and that is lost in stroma-dense tissue, where the same curves truncate and collapse toward random. The RSF index fell in a clean, near-monotonic progression, ∼450 (maximal-bias reference), ∼300, ∼150-160, ∼75, and ∼60 for the 30, 90, 180, and 320 µm gradients, and ∼40 for the random reference, mirroring the ordering of the curves themselves and resolving even the broad 180 and 320 µm gradients from each other and from random (Supplementary Fig. 4). Here the index and the curves tell the same story with the index distilling the ordering already visible in the curves into a single reproducible number per condition, its value being comparability across specimens rather than the recovery of distinctions the curves themselves have lost.

The **nearest-stroma distance distributions** themselves shifted and broadened in orderly fashion across conditions (Fig. 4g). The maximal-bias reference distribution rose steeply at short distances and fell off abruptly by ∼50-55 µm the most localized of the set; the 30 µm gradient peaked near 25 µm with a broad right-hand shoulder reaching ∼100-115 µm; and the 90 µm gradient gave a similarly defined but substantially broader profile that declined gradually across the measurable range, so that unlike in dense tissue the 30 and 90 µm conditions stayed clearly distinct. The 180 and 320 µm gradients overlapped more and drifted toward the flat random reference, yet their ordering and progressive broadening with interaction range remained visible; the measurable range extended to ∼175-200 µm, giving the distributions room to separate.

When condensed into summary statistics, this ordering became quantitative. The KS statistic spanned a broad, near-monotonic range of ∼0.8 (maximal-bias reference) to ∼0.45, 0.23, 0.12, and 0.08 for the 30, 90, 180, and 320 µm gradients, with low replicate variability. The **median** distance rose monotonically from ∼25 µm (maximal-bias reference) through the gradients to ∼65 µm (random), separating even the 180 and 320 µm scenarios. Together, these provide a sensitive readout for stroma-driven proliferation, as the KS statistic captures the full distributional shift from random to the maximal-bias reference, while the median is insensitive to the tails and replicate noise, making the pair well-suited to comparing specimens (Supplementary Fig. 4A(c,d)).

By contrast, the distance distribution’s **peak** location was unreliable. Distributions were multimodal or bimodal owing to lattice discretization; replicate variance was large, and the ordering across scenarios was non-monotonic and overlapping. The maximal-bias reference peak even shifted to ∼40 µm despite most cells lying much closer to the stroma. Because peak location depends on a single local maximum rather than on integrating the distribution, it does not provide a stable estimate of stromal spatial association, and we excluded it from all subsequent analyses (Supplementary Fig. 4A(b)).

The **RDF** likewise resolved both association strength and range here (Fig. 4i). The maximal-bias reference profile plateaued to ∼45-50 µm before declining to baseline near 150 µm, plateauing rather than peaking at a preferred radius because proliferating cells decorate the stromal boundary, so enrichment is greatest at the shortest distances and falls off only as the surrounding tumor compartment is reached. The gradient curves fell in order beneath it, each with a lower, shorter-reaching enrichment as its interaction range grew, down to the flat random profile. gmax ordered the conditions cleanly, ∼4.8 (maximal-bias reference), 3.5, 2.4, 1.5, 1.25 for the 30, 90, 180, and 320 µm gradients, and ∼1 for the random reference (Fig. 4i). The summary was most discriminating where enrichment was strong, spacing the near-stroma conditions widely, and less so among the long-range gradients, which sat closer together just above the random baseline; because it reads a single extremal point of a smoothed density ratio rather than integrating the curve, its replicate spread was the largest of the four summaries and grew as enrichment weakened. In this dispersed tissue geometry, then, geometric interference is minimal and the RDF recovers both the association strength and the relative range of stromal spatial association with high fidelity, gmax capturing that association strength faithfully where it is high and read with more caution as it approaches random.

The cross **J-function** gave the cleanest separation of any method (Fig. 4j). The maximal-bias reference curve fell to near zero by ∼45-50 µm, and the gradient curves declined in strict order, the 180 and 320 µm curves remaining distinguishable from each other, the strongest resolution of broad gradients observed in this study. **Jmin** traced the same ordering as a tight, monotonic progression, ∼0 (maximal-bias reference), ∼0.35-0.40, 0.75-0.80, 0.90, 0.95 for the 30, 90, 180, and 320 µm gradients, and ∼1 for the random reference (Fig. 4j). This summary was the most discriminating of the four. The adjacent conditions were separated by wide, even steps across the entire gradient range, so the broad 180 and 320 µm gradients, which the RSF index, median, and gmax still ordered but distinguished less sharply, remained clearly apart on the Jmin. Its replicate spread was also the smallest of any method, so those steps reflect the underlying gradient rather than sampling noise, and adjacent conditions do not overlap. In this favorable tissue geometry, the J-function therefore combines the range resolution of the curve with a summary that is both finely graded and highly reproducible, a pairing that makes it the most trustworthy discriminator of the four.

### Stromal architecture imposes a hard ceiling on the recoverable stroma-tumor cells spatial association range

The favorable behavior above is not guaranteed; it is a property of the tissue stromal architecture. In a high-content, high-dispersity specimen (1248 stromal pixels, edge fraction 0.81; Fig. 4a-e; Supplementary Fig. 3A), the characteristic spacing between neighboring stroma is only ∼75-80 µm. Beyond this distance, a tumor cell is, on average, closer to a different stromal structure, so separations larger than the inter-stromal spacing carry no information about a single source. Every method inherited this ceiling; the distance-dependent curves were effectively constrained to this short window, and the 180 and 320 µm gradients became difficult to distinguish despite arising from different imposed ranges. Mechanistically, once the assigned interaction range exceeds the inter-stromal spacing, most tumor cells encounter a nearer stromal structure before the imposed gradient has decayed, so the long-range signal is overwritten by the stromal architecture, and the observed distributions converge toward the random expectation.

The consequences for interpretation are central. In this high-content/high-dispersity specimen, the maximal-bias reference KS statistic was ∼0.75, whereas the exponential gradients compressed rapidly toward the random reference (Fig. 4b). The RDF similarly showed strong enrichment for maximal bias but substantial overlap among the broader gradients as they approached random (Fig. 4d). The RSF index separated maximal bias clearly from the other conditions, while the exponential gradients occupied a much narrower range near the random reference (Fig. 4c). Thus, a KS value, RSF index, or gmax has no universal absolute meaning since identical values from specimens with different stromal architectures can reflect different underlying spatial associations. The specimen-specific random and maximal-bias references define the interpretable dynamic range. Notably, even in this constrained regime, the cross J-function retained ordered separation across the reference states (Fig. 4e). The two low-dispersity regimes share the same geometry-dependent principle; Supplementary Fig. 5 shows additional point patterns and nearest-stroma distance summaries in Supplementary Figs. 6 and 7.

### Dispersity modulates signal association strength while content sets resolution

The two stromal architectures analyzed above and in the main figures are both highly dispersed. To separate the roles of stromal content and dispersity, we applied the same four methods to the two remaining regimes, high content with low dispersity and low content with low dispersity, in which the stroma form thick, bundled tracts rather than thin, dispersed ridges (Supplementary Fig. 5).

In the high-content, low-dispersity stromal architecture, the results mirrored the high-content, high-dispersity case in the main figures where the small inter-stromal spacing compressed the measurable range, so all four method curves truncated at a short distance, and the summary statistics collapsed toward the random reference. The maximal-bias reference KS ceiling was low, the median distance saturated quickly, the radial distribution function reached only modest enrichment, and the 180 and 320 µm gradients were barely separable from random. As elsewhere, the cross J-function retained the most residual separation among the broad gradients. The four gradients were poorly resolved. (Supplementary Fig. 5a-e; Supplementary Fig. 6 for point patterns and distance summaries).

In the low-content, low-dispersity stromal architecture, the results instead mirrored the favorable low-content, high-dispersity case, with the wide inter-stromal spacing restoring a long measurable range; the resource selection function and radial distribution function profiles were shifted outward and decayed at increasing distance as the gradient grew, and the cross J-function and nearest-stroma distance summaries progressed in monotonic, well-separated order from maximal-bias reference to random, with even the broad gradients distinguishable. The gradients were cleanly resolved. (Supplementary Fig. 5f-j; Supplementary Fig. 7 for point patterns and distance summaries).

Taken together, the four architectural regimes separate the roles of the two stromal architecture descriptors. Stromal content governs resolvability since both high-content stromal architecture, regardless of dispersity, compressed the gradients and capped the dynamic range at the short inter-stromal spacing, whereas both low-content stromal architecture resolved them out to long range, confirming that the recoverable spatial association range tracks stromal content rather than dispersity. Stromal dispersity instead appears to act on signal association strength through the accessible stromal surface, as the low-dispersity, bundled stromal architecture, which presents less tumor-contact interface, yielded summaries scaled down relative to their high-dispersity counterparts even where the ordering of conditions was preserved. *In short, stromal content decides whether the gradients can be distinguished at all, while stromal dispersity modulates how strong the recoverable signal is when they can*.

### The proliferative fraction independently changes every statistics

Stromal architecture is not the only confound. Holding stromal architecture fixed (∼650 stromal pixels and 0.53 stroma edge fraction, hence low dispersity), we varied the proliferative fraction across values that bracket the treatment course at 1%, 5%, 20%, 50%, and 75% (Fig. 5; Supplementary Figs. 8 and 9). Increasing the fraction pushes proliferating cells farther from stroma even when the underlying rule is unchanged, so only so many sites can lie adjacent to stroma; once filled, the remainder must be placed outward, broadening the spatial pattern and weakening the apparent signal. This occupancy effect compressed all metrics, but unequally. The KS statistic showed the strongest fraction dependence, with the maximal-bias reference decreasing from approximately 0.95-1.0 at 1% proliferation to ∼0.25 at 75% (Fig. 5b). The RSF index also decreased as proliferative fraction increased, compressing the separation among reference conditions (Fig. 5c). Jmin remained comparatively stable and showed the lowest replicate variability (Fig. 5d). The maximum RDF was the least reliable at low proliferative fractions. At 1% and 5%, sparse proliferating cells produced large replicate-to-replicate fluctuations in the local density estimate, whereas the metric became substantially more stable at 20% and above (Fig. 5e). The practical consequence is that identical statistics obtained at different proliferative fractions do not imply equivalent biology; proliferative fraction must therefore be accounted for alongside stromal architecture.

**Figure 5.**
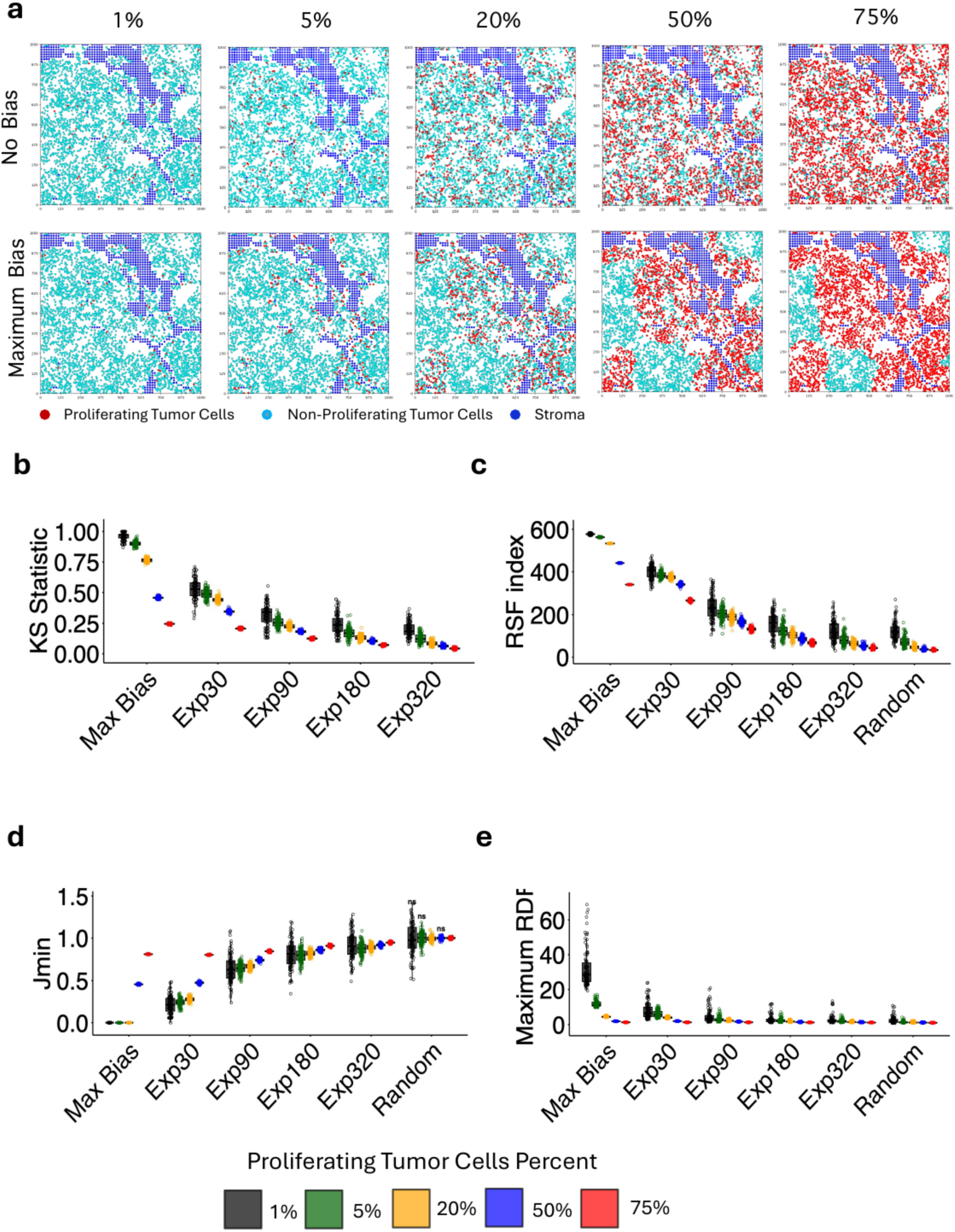
Proliferative fraction affects spatial-analysis accuracy. (a) Representative point patterns at 1%, 5%, 20%, 50%, and 75% proliferating tumor cells under no bias/random assignment (top) and maximum bias (bottom); proliferating tumor cells are red, non-proliferating tumor cells cyan, and stroma blue. (b-e) Summary statistics across the reference-state ladder (maximum bias, Exp30, Exp90, Exp180, Exp320, and random) for (b) KS statistic, (c) RSF index, (d) Jmin, and (e) maximum RDF. Colors denote proliferating-tumor-cell percentage as shown in the figure.

### The framework recovers a strong, short-range spatial association that departs from diffusion-only expectations in an experimental specimen

We applied the framework to an experimental specimen with low stromal content and high dispersity (304 stromal pixels, edge fraction 0.82), a favorable stromal architecture in which the analyses extend beyond 200 µm (Fig. 6a). To localize the specimen, we compared it against the reference states (Fig. 6) from maximal bias to the exponential gradients with characteristic distances of 30, 60, 90, and 320 µm, and the random/no-bias reference.

**Figure 6.**
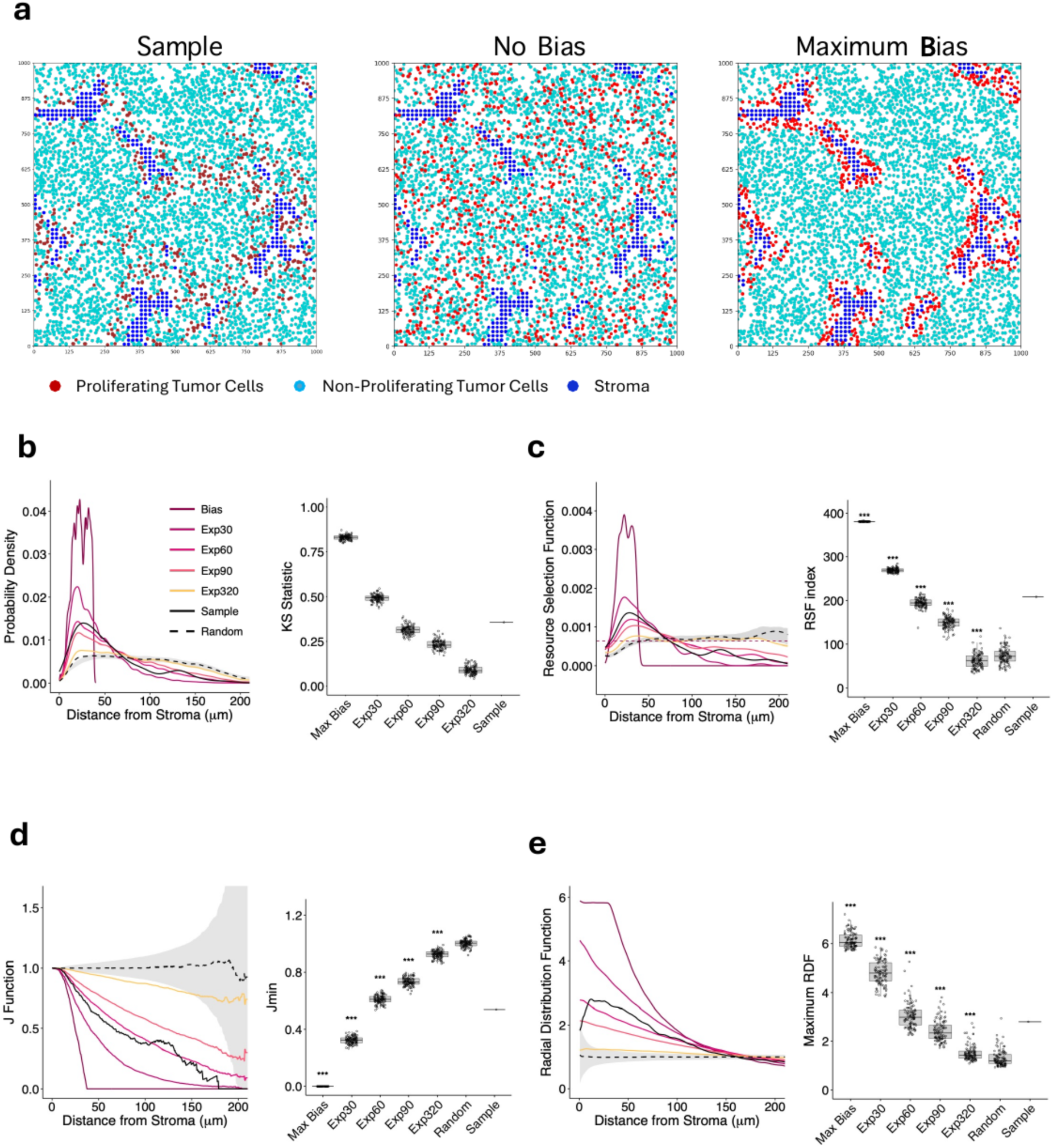
Application of the spatial statistical methods to a histological sample. An experimental ALK+ NSCLC sample is compared with in silico reference states generated on the same tissue geometry. (a) Point patterns for the experimental sample, no-bias/random reference, and maximum-bias reference; proliferating tumor cells are red, non-proliferating tumor cells cyan, and stroma blue. (b) Nearest-stroma distance probability density for maximum bias, Exp30, Exp60, Exp90, Exp320, random, and the experimental sample (left), with the KS statistic for the same conditions (right). (c) Resource selection function for the same reference states and sample (left), with RSF index (right). (d) Cross J-function (left) and Jmin (right). (e) Radial distribution function (left) and maximum RDF (right). The experimental sample is shown in black; the reference states are shown in the colors indicated in panel b.

All four methods agreed on a strong short-range stromal association. The nearest-stroma distance distribution for the experimental sample lay between the Exp30 and Exp60 references, and its KS statistic was ∼0.36-0.38, between the corresponding reference values and well above the random reference (Fig. 6b). The RSF likewise placed the sample between the short-range Exp30 and Exp60 references, while the broader reference gradients approached the random baseline (Fig. 6c). The cross J-function for the sample showed the same ordering with the sample trace laying between the short-range reference curves over much of the measured interval, and the sample Jmin being ∼0.55-0.60, compared with ∼0 for maximal bias and ∼1 for the random reference (Fig. 6d).

The RDF was particularly informative (Fig. 6e). Rather than matching a single exponential reference throughout its full shape, the experimental sample showed a pronounced short-range peak followed by a broader decay, with maximum RDF ∼2.8. This was lower than the maximal-bias reference (∼6) but clearly above the random reference (∼1-1.2). The mismatch between the experimental RDF shape and any single exponential reference suggests that a simple exponential distance-decay pattern does not fully describe the observed stromal-associated proliferation and may reflect additional biological organization, such as threshold responses to stromal signals, extracellular-matrix structure, or heterogeneous stromal subpopulations.

Taken together, all four methods place this specimen at a strong, short-range stromal spatial association of ∼30-60 µm, roughly two to four tumor-cell diameters, while the systematic departures from the theoretical gradients indicate that stromal-associated proliferation is not fully explained by simple diffusion. The key check is that the four methods, which fail under different conditions, converge on the same range and on the same conclusion of strong-but-incomplete spatial association. This makes the result robust to the weaknesses of any single statistic and shows that the calibrated framework can extract an interpretable interaction range from a real specimen rather than only from controlled simulations.

## Discussion

Tumor function is inherently based on spatial associations between tumor cells and their tissue environment. A tumor cell’s behavior is shaped not only by its own state but by the local neighborhood, specifically which cells and structures lie nearby and how close. Histology captures this organization directly, yet quantifying it as whole-section counts, densities, and “fraction-positive” summaries collapses spatially structured tissue into a single number, averaging over the local relationships that drive processes such as immune exclusion, metabolic support, and treatment resistance. Two specimens with identical positive fractions can differ completely in how those positive cells are arranged relative to the structures that matter (Fig. 1a,b). Recovering that arrangement quantitatively and reproducibly requires spatial statistics rather than bulk quantification.

We developed a framework that measures stromal-associated proliferation directly from histology by calibrating four complementary spatial statistics against biologically grounded reference states, and evaluated it across stromal architecture, proliferative fraction, and simulated spatial association strength. It addresses the three questions posed at the outset whether a spatial association is *present* (deviation from the random reference), the *range* over which it acts (where the RSF, J-function, and RDF return to baseline), and its *association strength* (the position of each statistic within the random-to-maximal-bias reference interval).

The main message is that two tissue properties, not the analytical method, set the limits of what can be learned; specifically, stromal architecture caps the recoverable range through inter-stromal spacing, and proliferative fraction independently compresses every metric through occupancy. Specifically, stromal content, through the inter-stromal spacing it sets, governs whether the gradients can be resolved at all, while stromal dispersity, through the accessible tumor-contact surface, modulates the association strength of the recoverable signal once content permits it. Content therefore decides resolvability and dispersity decides association strength, so a specimen’s two stromal architecture descriptors have to be read together rather than collapsed into a single notion of “dense” versus “sparse” stroma. Because both act on the observable signal rather than the biology, spatial statistics from tissue must be read relative to specimen-specific reference states and proliferative fraction rather than on absolute scales the random and maximal-bias reference models supply exactly this calibration.

The framework’s practical value lies in how to interpret each summary statistic when tissue geometry and proliferative fraction vary, since both act on the statistic rather than the biology. The RSF index measures the integrated departure of proliferative intensity from the random baseline; stromal geometry can compress the distance window over which it is informative, while increasing proliferative fraction reduces separation among reference conditions. The median nearest-stroma distance is reproducible but can saturate once the interaction range exceeds the geometry-imposed measurable window, and it shifts with proliferative fraction even when the underlying assignment rule is fixed. The KS statistic is highly sensitive to spatial bias, but its dynamic range and separation among gradients depend on stromal architecture and proliferative fraction, so interpret it relative to specimen-specific references rather than an absolute threshold. Jmin (the J value at 60 µm used here) is comparatively stable across replicates, although reducing the full J curve to a single value necessarily loses information about curve shape and long-range separation. The RDF maximum, gmax, provides a direct measure of local enrichment but is especially sensitive to sparse proliferating-cell counts and is unreliable at the lowest proliferative fractions. Across all four summaries, the specimen-specific random and maximal-bias references provide the appropriate calibration.

This yields a tiered strategy. Jmin (the J-function value at 60 µm) and the median distance are the reproducible readouts for screening many specimens; the RSF index and gmax quantify association strength, the latter only where proliferation is sufficient; the full RSF, J, and RDF curves resolve range and mechanism in well-powered samples; and peak location is avoided throughout. The overarching rule is that no summary statistic carries meaning on its own; each is read as a position within the specimen-specific random-to-maximal-bias reference interval, at that specimen’s proliferative fraction, and agreement among statistics that fail under different confounds remains the strongest evidence of a real spatial association.

Several limitations bound these conclusions. The analysis is performed on two-dimensional sections, so inferred distances are projections of a three-dimensional tissue geometry. Discretizing the stroma onto a ∼15 µm lattice sets a resolution floor and is the source of the multimodality that renders peak location unusable. Because every scalar is interpreted within a specimen-specific random-to-maximal-bias reference interval, values cannot be pooled across specimens without recomputing the reference states, and the maximal-bias reference ceiling itself shifts with proliferative fraction. The framework quantifies spatial association but does not by itself establish mechanism, so the departure of the observed RDF from the diffusion gradients is suggestive of threshold or matrix spatial associations rather than proof of them and would require orthogonal validation. Finally, the experimental demonstration rests on a single favorable specimen; extending the calibration across a full treatment time course and a specimen cohort remains future work.

Applied to tissue, the framework identified strong stromal-associated proliferation over ∼30-60 µm, with systematic deviations from diffusion-only gradients that implicate threshold signaling, matrix structure, or stromal heterogeneity. Because it depends only on a segmented point pattern and a definition of the reference states, the approach is not specific to stroma or proliferation, and any pairing of a spatial covariate with a labeled population immune infiltrate relative to vasculature, hypoxia relative to necrosis can be analyzed in the same calibrated way, and across a treatment time course, where Stromal architecture and proliferative fraction both shift, that specimen-specific calibration is what makes comparisons meaningful.

## Methods

### Tumor model, treatment, and tissue collection

Xenograft studies were performed by bilateral subcutaneous implantation of 5 × 10^6^ tumor cells per injection suspended in 100 µL of a 1:1 mixture of RPMI and Basement Membrane Extract (BME) Type 3 (R&D Systems, #363201002P) into 4-6 -week-old NOD.Cg-Hgftm1.1(HGF)Aveo Prkdcscid Il2rgtm1Wjl/J mice (NSG-hHGF mice) of both sexes. The recipient animals were bred in-house from breeders obtained from Jackson Laboratory, Strain #014553. Treatment was initiated 3 weeks after tumor implantation. Small-molecule inhibitors were administered once daily by oral gavage (100 µL per dose). Alectinib (20 mg/kg) was dissolved in water. Tumor dimensions were measured weekly using electronic calipers, and tumor volumes were calculated assuming spherical tumor tissue geometry. At the study endpoint, tumors were excised, weighed, and fixed in 10% neutral buffered formalin. All animals were euthanized before tumors reached the threshold size of 2,000 mm^3^ specified in the animal protocol. 30-45 minutes prior to euthanasia, the animals were injected IP with 10 mg/ml BrdU dissolved in 1X PBS. All xenograft studies were conducted in accordance with the approved Institutional Animal Care and Use Committee (IACUC) protocol (#IS00005557) at H. Lee Moffitt Cancer Center. Animals were housed in an AAALAC-accredited, specific pathogen-free vivarium under veterinary supervision and maintained under standard conditions of temperature and humidity with a 12-hour light/12-hour dark cycle. Immunohistochemistry and image segmentation *were performed as described in*^28^. Tissue segmentation was performed using the AI-assisted digital pathology platform Aiforia version 5.2 as described in^38^. Tissues were segmented into BrdU+/- tumor cells, stroma, and necrotic tissue. We segmented stromal areas identified by counterstaining as polygons separating cellular and acellular ECM components. Whole-slide images were segmented with Aiforia into three tissue classes: BrdU+ and BrdU-tumor cells, and stroma, the latter comprising fibroblasts and extracellular matrix rather than a single cell type (Fig. 1d; Supplementary Fig. 1).

### Point-pattern construction

Segmented images were converted into marked spatial point patterns within an observation window defined by the tissue boundary. Each tumor cell contributed one point at its nuclear centroid carrying a mark of proliferative status (BrdU+ or BrdU−). Stroma, a continuous composite scaffold rather than discrete cells, was discretized into points on a regular square grid of spacing ≈15 µm (about one tumor-cell diameter), so that stromal abundance, tissue geometry, and boundary length were preserved at single-cell resolution. The stromal edge, the contour separating stromal and tumor regions, was the set of stromal grid points immediately adjacent to tumor tissue. For every tumor cell, the distance to the nearest stromal point was computed and used as the fundamental spatial covariate throughout. All spatial computations were performed in R with the spatstat package, which supplies the point-pattern representation, spatial-covariate handling, edge-corrected summary functions, and Monte-Carlo envelopes used below.

### Random reference and reference states

We defined a random reference to preserve tissue architecture while removing any preferential spatial association between proliferation and stroma. BrdU+ cells were held exactly as observed, and proliferative status was permuted at random across the tumor-cell population (Fig. 3a). This preserves tissue architecture, stromal architecture, tumor-cell density, and proliferative fraction while removing any spatial association of proliferation with stromal proximity, defining the lower bound of the achievable signal (random reference). Because stromal architecture remodels substantially during treatment, preserving the observed stromal architecture within each specimen is what isolates proliferative localization from broader treatment-induced changes in tissue structure.

As the matching upper bound, we constructed a maximal-bias reference in which we assigned proliferative status preferentially to the tumor cells closest to stroma, decorating the stromal network with proliferating cells, the strongest stromal spatial association the specimen can physically express. Its interaction range depends on both stromal architecture and proliferative fraction, so when stroma are scarce or proliferating cells are abundant, even maximal-bias reference must place cells progressively farther from stroma, so the maximal-bias reference distribution can extend to 60-120 µm and a short-range exponential gradient (e.g. *μ* = 30 µm) can be more localized than maximal-bias reference itself. The maximal-bias reference therefore measures the stromal carrying capacity for proliferation in a given specimen rather than a fixed distance. In low-content specimens, this can reverse the expected ordering of the distance-based summary statistics, the median and peak location between maximal-bias reference and the 30 µm gradient, so those summaries are interpretable only relative to the maximal-bias reference computed for the same specimen. Each summary statistic was interpreted by its position within this specimen-specific random-to-maximal-bias reference interval rather than on an absolute scale, because the maximum expressible signal is fixed by tissue stromal architecture.

### Controlled stromal-spatial association gradients

To benchmark method sensitivity against a known ground truth, we generated synthetic proliferative patterns with prescribed range and association strength while preserving the observed tissue Stromal architecture and proliferative fraction (Fig. 3b,c). The biological rationale is that stroma-derived paracrine factors diffuse while being consumed and degraded, so their concentration and the proliferation probability they support decline approximately exponentially with distance from the stroma. Accordingly, for each tumor cell, the nearest-stroma distance *r* was mapped to a proliferation probability through an exponential cumulative distribution function,

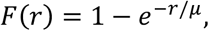

in which the interaction range *μ* sets the effective interaction range; a fraction 1 − *e*^−1^ ≈ 0.632 of proliferating cells falls within *μ* of stroma. Proliferative status was assigned by sampling tumor cells in proportion to this probability until the observed BrdU+ count was reached, holding the proliferative fraction fixed. Gradients spanned *μ* ≈ 15-320 µm (1-21 tumor-cell diameters). Together with the random reference and the maximal-bias reference upper bound, these gradients establish a continuous spectrum of stromal spatial associations from random reference to maximal stromal spatial association. Applied to each statistic in turn, the null fixes its zero, maximal-bias reference its specimen-specific maximum, and the gradients supply known-range ground truth, so that the sensitivity, specificity, and resolution of every statistic and the conditions under which each fails were evaluated against the same benchmark across tissue Stromal architecture and proliferative fraction rather than assumed.

### Resource selection function

We quantified how proliferative intensity varies with proximity to stroma using a resource selection function, adapted from ecology, where it expresses the relative probability that a unit of space is used as a function of its environmental covariates. An inhomogeneous point process with intensity *λ*(*u*) at location *u* is modeled as a function of a spatial covariate *Z*(*u*)^23,34^,

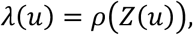

where *ρ*(·) is the relative-intensity function and *Z*(*u*) is the distance from *u* to the nearest stroma. Rather than assuming a parametric form, *ρ* was estimated nonparametrically by kernel-smoothed regression of point intensity on the covariate (the rhohat estimator in spatstat), effectively tessellating the tissue into bands at increasing distance from stroma and estimating proliferative intensity in each. A value of *ρ*(*z*) above baseline indicates that proliferating cells occur more frequently at distance *z* than expected from spatial availability. The estimate was interpreted relative to the random-assignment control rather than the tissue-average intensity, because proliferating cells are a subset of all tumor cells and only the random reference carries the appropriate baseline.

Two choices govern the estimate *ρ*(*z*). The first is whether to correct for tumor-cell availability. Left as an intensity (proliferating cells per unit area), ρ(z) is confounded by the uneven distribution of tumor cells around stroma; dividing instead by the local density of all tumor cells gives the fraction of tumor cells that are BrdU+ at distance z, a quantity that is flat at the overall proliferative fraction when labels are assigned at random. The second concern concerns smoothing. The grid and bandwidth can be chosen automatically for each pattern or fixed to a common grid and a bandwidth of one cell diameter (about 15 µm) applied in the same way to the observed pattern and to every simulation. Together these choices produce four variants (v1, no baseline and floating bandwidth; v2, baseline and floating bandwidth; v3, no baseline and fixed bandwidth; v4, baseline and fixed bandwidth). We report v4 throughout unless stated otherwise, because it corrects for availability and uses a prior bandwidth, which together make the random-labeling envelope test valid;

For comparison across conditions, we summarized each estimated curve by a single scalar, the RSF index, the integrated absolute departure of the relative intensity from its baseline,

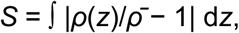

where *ρ*(*z*) is the kernel-smoothed relative intensity at distance *z* from stroma and *ρ̄* is the average intensity, i.e. the constant value expected under no distance spatial association. *S* measures the total association strength of distance-spatial association selection, integrating enrichment and depletion across the full distance range, and is reported in the units of the integrated curve. Because it averages over the whole curve, *S* is more reproducible across replicates than the RSF peak height or peak location, which are sensitive to the smoothing bandwidth; it therefore serves as an association strength summary complementary to the KS statistic, which tests the significance of the covariate spatial association. The random-assignment control yields a nonzero *S* owing to finite-sampling fluctuation of the estimate, so each condition’s index is interpreted relative to that floor rather than against zero. The four variants are compared in Supplementary Note 4, Supplementary Table 1, and Supplementary Figs. 3B(d-f) and 4B(d-f).

### Cross J-function

The multitype (cross) J-function compares the observed proximity of one point type to another against the proximity expected from availability. For types *i* (proliferating tumor cells) and *j* (stroma)^23,36^,

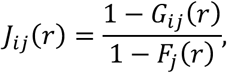

where *G*ij(*r*) is the cross nearest-neighbor distribution, the probability that the nearest stroma lies within *r* of a typical proliferating cell, and *F*j(*r*) is the empty-space function, the probability that the nearest stroma lies within *r* of an arbitrary location, i.e., stromal availability across the tissue. Under independence, *J*ij(*r*) ≡ 1; *J*ij(*r*) < 1 indicates that proliferating cells lie closer to stroma than expected (positive spatial association), and the distance at which *J*ij(*r*) approaches zero estimates the range within which essentially all proliferating cells lie near stroma. We used edge-corrected estimators of *G*ij and *F*j (Jcross in spatstat) and summarized each curve by its value at 60 µm (∼four tumor-cell diameters), near the upper end of the biologically relevant range while remaining within the measurable window.

### Radial distribution function

The radial distribution function (RDF; equivalently the pair correlation function) describes the relative density of point pairs as a function of their separation. In two dimensions, it derives from Ripley’s K-function^23^,

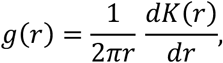

and for two types, generalizes to the cross RDF *g_i_*_j_(*r*) = *ρ*(2)(*r*)/5*λ_i_λ*_j_7, with *ρ*(2)ij the cross-product density and *λ* the intensities. Under independence *g*ij(*r*) ≡ 1; values above one indicate enrichment of proliferating cells at distance *r* from stroma, values below one indicate depletion. We estimated *g*ij(*r*) between proliferating tumor cells and stroma with edge correction and kernel smoothing (pcfcross in spatstat) and summarized each curve by its maximum, *g*max. Because kernel estimates of a density ratio from a few points are noisy, *g*max is sensitive to sampling and can exceed 1 even under randomness, so it was interpreted alongside the random reference and treated as unreliable below ∼20% proliferation.

### Nearest-stroma distance distributions

For each tumor cell the distance to the nearest stromal point was recorded, generating separate nearest-stroma distance distributions for BrdU+ and BrdU-cells and for each simulated condition. Distributions were summarized by their median (a robust central tendency) and by a two-sample Kolmogorov-Smirnov (KS) statistic against the random (or non-proliferating) control^33^,

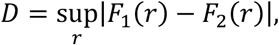

the maximum vertical distance between the two empirical cumulative distribution functions, which is sensitive to differences across the entire distribution rather than a single moment. Smaller medians and larger KS values indicate stronger stromal proximity. Peak location proved unstable, multimodal due to lattice discretization, with large replicate variance and non-monotonic ordering, so we excluded it from quantitative interpretation.

### Stromal architecture descriptors

Each specimen was characterized by two descriptors. Stromal content is the total number of stromal grid points, a measure of overall abundance. Stromal edge fraction is the proportion of stromal points lying on the stromal-tumor boundary relative to total content, a measure of dispersity and accessible signaling surface. This way, values near 1 indicate thin, dispersed stroma in which nearly all stroma contacts tumor, whereas low values indicate thick, bundled stroma with large interior regions not in contact with tumor (Supplementary Fig. 10). Crossing high versus low content with high versus low edge fraction defines the four architectural regimes analyzed (Fig. 4; Supplementary Fig. 5).

### Replication, confidence envelopes, and comparison

The null, the maximal-bias reference, and every exponential gradient were generated in 99 independent stochastic realizations; reported curves and summary statistics are the mean across realizations, with replicate spread shown where relevant. Pointwise confidence envelopes were constructed from repeated realizations of the random reference and used to distinguish biologically meaningful deviations from variation expected under tissue Stromal architecture alone. All spatial computations, point-pattern handling, the distance-to-stroma covariate, edge-corrected summary functions (rhohat, Jcross, pcfcross, and the nearest-neighbor and empty-space functions), and simulation envelopes were performed in R with spatstat.

### Classical spatial methods for comparison

For comparison, we also applied conventional spatial statistics: quadrat-count correlations, Ripley’s cross-K function, the Morisita index, and kernel density colocalization, each of which requires the interaction range to be fixed in advance through a quadrat size or smoothing bandwidth. Their scale-dependent spatial association and resulting limitations in this setting are detailed in Supplementary Note 1 and Supplementary Fig. 2.

## Data availability

https://github.com/tmiti26/SpatialStatMethods

## Code availability

https://github.com/tmiti26/SpatialStatMethods

## Acknowledgements

This project was supported by pilot grants from Moffitt Cancer Center Evolutionary Therapy Center of Excellence (to AM and DB) and Lung Cancer Center of Excellence (to AM), DOD LC190085 (to AM and DB). This work has been supported in part by the Analytic Microscopy Core, Vivarium Core, and the Tissue Core at the H. Lee Moffitt Cancer Center & Research Institute, a comprehensive cancer center designated by the NCI. TM and DB thank Dr. Chandler Gatenbee in the IMO Department at Moffitt for constructive criticism of the manuscript, and Dr. Anna Gaffney (Miller) and Dr. Philipp Altrock for contributions to the start of this project.

## Author contributions

TM conceived the analytical framework, selected and combined the spatial-statistical methods, performed all spatial analyses, prepared the figures and wrote the manuscript. BD and AM carried out the experiments. DM performed the image segmentation and contributed to figure preparation. AM and DB contributed to the development of the analytical approach and revised the manuscript. Claude (Anthropic) and ChatGPT were used to assist with analysis code (automated part of the code), figure preparation (made sure the figure style is the same across all figures), and editing of the manuscript text (checked for correctness of referred figures in the text and formatted for bioRxiv). We used Grammarly to check grammar and sentence structure. The authors reviewed all output and take full responsibility for the content. For transparency, early versions of this manuscript and code can be found on the GitHub repository as well.

