## supplementary information for "A reference-calibrated spatial-statistical framework for quantifying stroma-tumor cell spatial associations from histology and its geometric limits"

<sup>3</sup> TBD

<sup>4</sup> Department of Molecular Medicine, University of South Florida, Tampa, Florida.

#### **Contents**

Supplementary Note 1. Limitations of classical spatial statistical methods

Supplementary Note 2. Stromal architecture across the four content and dispersity regimes

Supplementary Note 3. Proliferative fraction in additional stromal architectures

Supplementary Note 4. Four estimators of the resource selection function

Supplementary Table 1. Estimators of the resource selection function

Supplementary Figures 1 to 10, with legends

### Supplementary Note 1. Limitations of classical spatial statistical methods

Before developing the reference-calibrated framework presented in the main text, we applied four established spatial statistics to the segmented point patterns: quadrat counts, Ripley's K-function, the Morisita index, and kernel density estimation. Each provided a useful exploratory description but could not answer the central question, because each requires the spatial scale of the association to be fixed in advance rather than recovered from the data (Supplementary Fig. 2).

**Quadrat counts.** Partitioning the tissue into quadrats and correlating per-quadrat stromal content with proliferative intensity detects coarse co-variation, but the result depends heavily on the chosen quadrat size and position, and it does not measure the relationship between tumor cells and stroma directly. In our data, the inferred correlation was not even stable in sign, as reducing the quadrat side from 700 $\mu\text{m}$ to 500 $\mu\text{m}$ was sufficient to change the correlation between stromal content and proliferation from positive ($R \approx 0.48$) to negative (Supplementary Fig. 2a). A method whose conclusion reverses with an arbitrary choice of scale cannot establish the existence or range of a stromal association when that scale is the unknown of interest.

**Ripley's K-function.** The cross-K function quantifies association between proliferating cells and stroma, but it accumulates that association across all distances up to a given radius, so a departure from spatial randomness signals an association without localizing it (Supplementary Fig. 2b). Because K characterizes global structure rather than local intensity, it also cannot separate genuine stromal proximity from the two components merely occupying overlapping regions of a heterogeneous tissue.

**Morisita index.** The Morisita index summarizes clustering of proliferating cells without any reference to stromal location, so an elevated value is compatible with many mechanisms unrelated to stroma (Supplementary Fig. 2c). Its characteristic scale conflates clustering parallel and perpendicular to stroma and, in branching stromal networks, cannot be translated into a biologically meaningful distance of stromal association. The index is also scale-dependent, decaying toward one as the quadrat diameter grows.

**Kernel density estimation.** Kernel-smoothed density maps render stromal and proliferative distributions continuously and can suggest colocalization, but the apparent structure depends entirely on the smoothing bandwidth (Supplementary Fig. 2d), since narrow bandwidths produce localized peaks that follow the stroma, while modest increases smooth them away and weaken the apparent association. The bandwidth is an externally imposed parameter rather than a biologically derived quantity, and the maps remain descriptive, yielding neither a range nor a strength of association.

Together, these methods can reveal that spatial organization exists, but they take the scale of association as an input rather than returning it as an output. This motivated the reference-calibrated framework of the main text, in which the range and strength of stromal association emerge from the data relative to specimen-specific random and maximal-bias reference states.

### Supplementary Note 2. Stromal architecture across the four content and dispersity regimes

The main text presents the two high-dispersity stromal architectures (high and low stromal content) in Fig. 4. The supplementary figures extend that analysis in three ways.

First, Supplementary Figs. 3A and 4A show, for the high-content/high-dispersity and low-content/high-dispersity specimens of Fig. 4, the point patterns for every reference state (maximal bias, exponential gradients of 30, 90, 180 and 320 $\mu\text{m}$, and random) together with the three scalar summaries of the nearest-stroma distance distribution, specifically peak location, KS statistic and median. The peak-location panels show directly why this metric was excluded from quantitative interpretation in the main text. Its replicate spread is large relative to the separation between neighboring conditions, so adjacent reference states overlap, whereas the KS statistic and the median computed from the same distributions separate the conditions with little replicate variation (Supplementary Figs. 3A, 4A, 6 and 7). The KS statistic is calculated using two reference populations; the main figures compare BrdU+ cells with randomly assigned BrdU+ labels, whereas Supplementary Figs. 3A, 4A, 6 and 7 compare BrdU+ cells with BrdU- tumor cells.

Second, Supplementary Figs. 3B and 4B give the global-envelope versions of the curve statistics for the same two specimens, each with the global $p$-value per reference state. In the low-content/high-dispersity specimen (Supplementary Fig. 4B), every gradient departs from the random reference on every statistic ($p = 0.005$, except $p = 0.020$ for the 320 $\mu\text{m}$ gradient on the radial distribution function), while the random realizations themselves are not significant ($p = 0.43$ to $0.56$). In the high-content/high-dispersity specimen (Supplementary Fig. 3B), the maximal-bias reference and the 30 $\mu\text{m}$ gradient remain significant on every statistic ($p = 0.005$), and the 90 $\mu\text{m}$ gradient on all but the radial distribution function ($p = 0.105$), but the 180 and 320 $\mu\text{m}$ gradients cannot be distinguished from random on any statistic ($p = 0.055$ to $0.595$). The whole-curve tests therefore reproduce the geometric ceiling described as in the main text, where once the imposed range exceeds the inter-stromal spacing, the gradient is no longer detectable.

Third, Supplementary Figs. 5 to 7 cover the two remaining regimes, high content with low dispersity and low content with low dispersity, in which the stroma forms thick, bundled tracts rather than thin, dispersed ridges. Supplementary Fig. 5 shows the full set of method curves and scalar summaries in the layout of Fig. 4, and Supplementary Figs. 6 and 7 show the corresponding point patterns and distance summaries. As described in the main text, the high-content/low-dispersity specimen behaves like its high-dispersity counterpart, with a short measurable window (under 100 $\mu\text{m}$) and long-range gradients that crowd toward the random reference, whereas the low-content/low-dispersity specimen resolves the gradients in order out to about 200 $\mu\text{m}$. The stromal edge fraction used to define dispersity is illustrated in Supplementary Fig. 10.

#### **Supplementary Note 3. Proliferative fraction in additional stromal architectures**

To test whether the occupancy-driven compression of the summary statistics is specific to the stromal architecture used in Fig. 5, we repeated the proliferative-fraction series (1%, 5%, 20%, 50% and 75% proliferating tumor cells) on two further stromal architectures (Supplementary Figs. 8 and 9). The same behavior was observed in both. For the exponential gradients, the KS statistic and the RSF index decreased steadily as the proliferative fraction increased, so the separation among reference states narrowed at high fractions; for the maximal-bias reference, they changed less, with a clear drop only at 75% in Supplementary Fig. 8. The maximum RDF was highest and most variable at 1% and 5%, where few proliferating cells enter the density estimate, and converged toward 1 at higher fractions. $J_{\min}$ changed comparatively little with proliferative fraction, although its replicate spread was widest at 1%. The occupancy effect is therefore a general geometric consequence of the finite number of tumor-cell positions near stroma rather than a property of a single stromal architecture.

### Supplementary Note 4. Four estimators of the resource selection function

The resource selection function $\rho(z)$ measures how the propensity for proliferation changes with the distance $z$ to the nearest stroma. Two independent choices in its estimation, one about availability and one about smoothing, give four variants that answer slightly different questions and that differ in whether they support a formal test.

**Availability correction (baseline).** Proliferation happens only where tumor cells are, and tumor cells are not evenly distributed relative to stroma. Without a baseline, $\rho(z)$ is an intensity in cells per unit area; with random labels, it doesn't come out flat but follows the tumor-cell availability profile, so both the curve and the RSF index report where cells are rather than where they proliferate. Dividing by the local density of all tumor cells corrects this. $\rho(z)$ then becomes a fraction between 0 and 1; it sits flat at the overall proliferative fraction under random labeling, and any departure from that flat line reflects genuine stroma-associated selection.

**Grid and bandwidth.** With a floating bandwidth, both the grid and the amount of smoothing are read off each pattern. The bandwidth then depends on how many cells there are and how they are arranged, so the observed curve and each random-labeling simulation are smoothed by different amounts, and smoothing itself becomes a confounder in the comparison; the estimate also varies more from one specimen to the next. Fixing the grid and setting the bandwidth to one cell diameter (about 15 $\mu\text{m}$) for the observed curve and every simulation removes this problem, because any difference that remains must come from the spatial arrangement of proliferation rather than from smoothing. A fixed bandwidth is also reproducible across specimens.

The four variants and their trade-offs are summarized in Supplementary Table 1.

**Supplementary Table 1. Estimators of the resource selection function.** Variants are defined by availability correction (rows) and by the choice of bandwidth and grid (columns).

| | Floating bandwidth, no grid | Fixed bandwidth and grid (about 15 $\mu\text{m}$) |
| --- | --- | --- |
| **No baseline (intensity per area)** | **v1.** Descriptive intensity curve. It carries both the availability and the smoothing confounders and cannot support a formal test. | **v3.** Supports a valid envelope test because the smoothing is common to all curves, and the bandwidth is reproducible, but the curve and the RSF index still reflect availability, so the random reference is not flat. |
| **Baseline (availability-corrected)** | **v2.** Corrects for availability, so the random reference is flat, but the bandwidth still varies between curves and the envelope test is therefore not strictly valid. | **v4 (used for inference).** Corrects for availability and supports a valid test, resulting in a resource selection function estimated with a fixed, prior bandwidth. |

In short, v1 needs no tuning but carries both confounders and cannot support a test. v2 clears the availability confounder while leaving the smoothing one. v3 supports a valid test but still mixes in availability. v4 removes both and is the version used for inference in the main text. Its only added cost is a second, spatial smoothing parameter for the baseline density, which we set to one cell diameter so that it matches the bandwidth used for the covariate.

Supplementary Figs. 3B and 4B (panels d to f) show the resource selection function of the two specimens of Fig. 4 under three estimation settings, given in the panel titles (bw, smoothing bandwidth; grid, evaluation grid), specifically, no fixed bandwidth and no grid (panel d, corresponding to v1), fixed bandwidth and fixed grid (panel e, corresponding to v3), and fixed bandwidth and fixed grid with normalization (panel f).

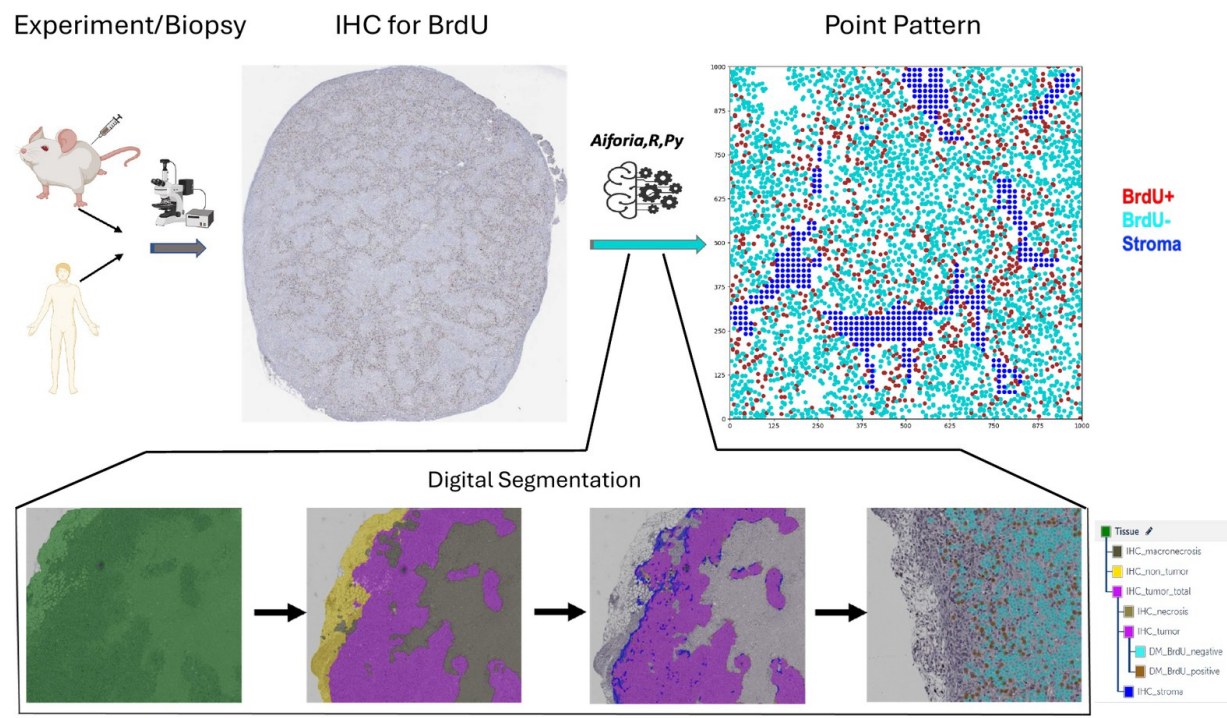

**Supplementary Fig. 1. From histological specimen to marked point pattern.** Workflow for converting an experimental or biopsy specimen into the input of the spatial analysis. Top: the specimen is sectioned and stained for BrdU by immunohistochemistry (IHC), and the whole-slide image is processed (Aiforia, R, and Python) into a marked point pattern in which BrdU+ tumor cells are red, BrdU- tumor cells cyan, and stroma blue. Bottom: successive layers of the digital segmentation, from tissue detection, through separation of tumor, stroma, and necrosis, to the classification of individual BrdU+ and BrdU- tumor cells; the hierarchy of segmentation classes is shown at right.

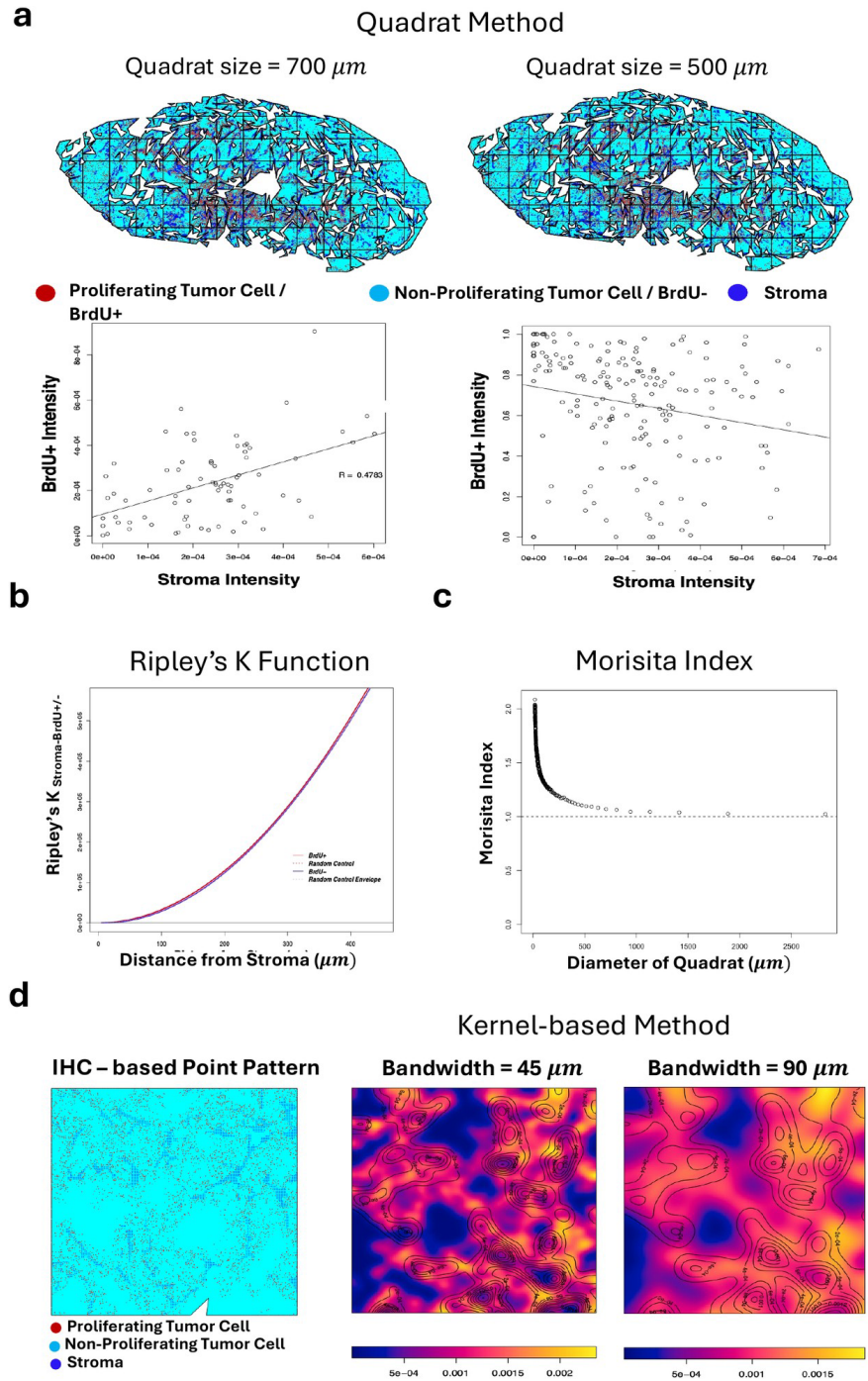

**Supplementary Fig. 2. Classical spatial methods have a limited ability to quantify stroma-tumor cell spatial associations.** (a) Quadrat method. Per-quadrat BrdU+ intensity plotted against stromal intensity at quadrat sizes of 700 $\mu\text{m}$ and 500 $\mu\text{m}$; the correlation changes from positive ($R \approx 0.48$) to negative as the quadrat size changes, showing dependence on an arbitrary scale. (b) Ripley's cross-K function as a function of distance from stroma. (c) Morisita index as a function of quadrat diameter, decaying toward 1 and therefore scale-dependent. (d) Kernel-based density maps computed from an IHC-derived point pattern at bandwidths of 45 $\mu\text{m}$ and 90 $\mu\text{m}$, showing that the inferred density structure depends on the chosen bandwidth. Together, the panels show that classical methods require the scale of association to be fixed in advance.

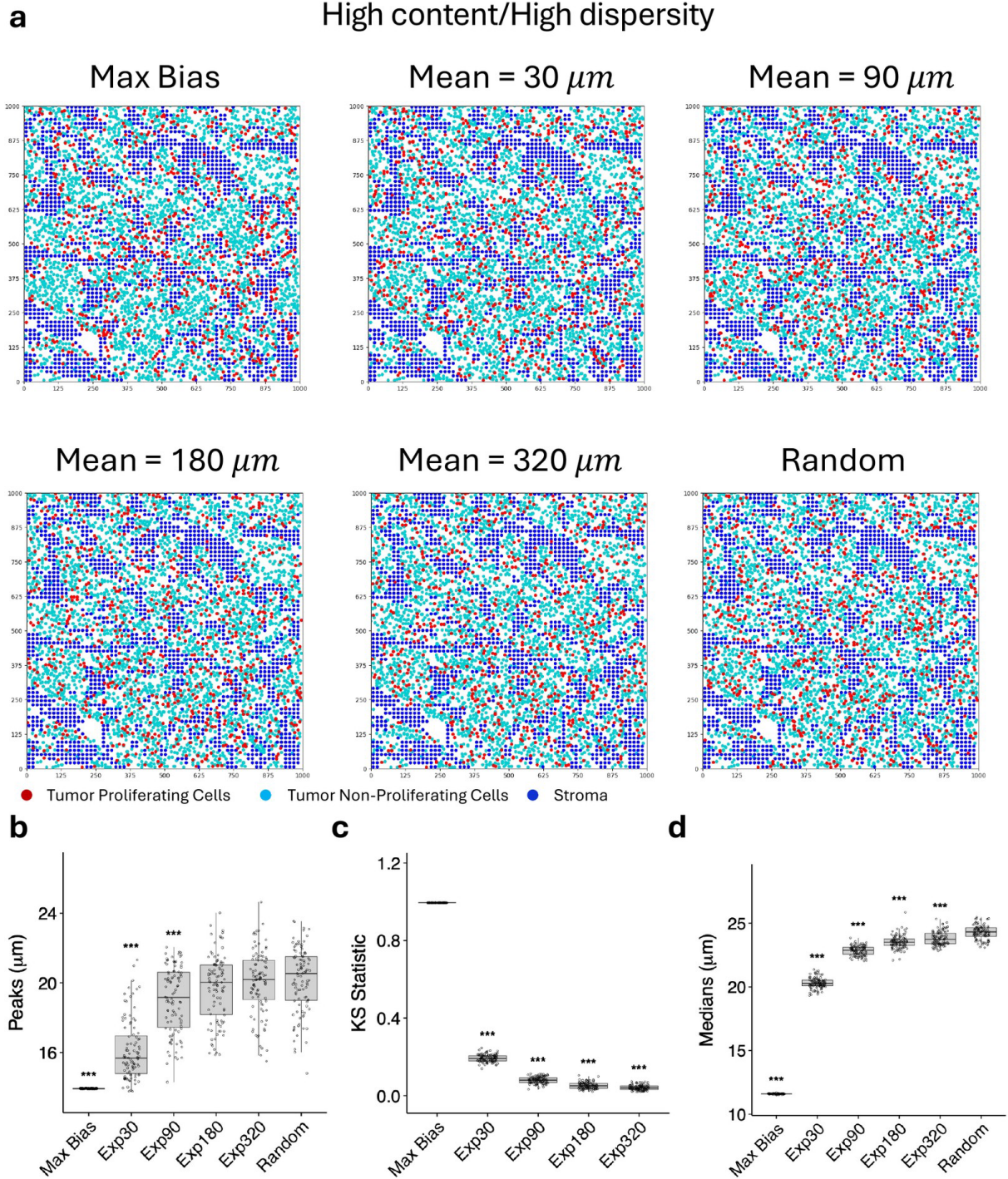

**Supplementary Fig. 3A. High stromal content and high dispersity: point patterns and distance summaries.** Additional analysis of the high-content/high-dispersity specimen of Fig. 4a-e. (a) Point patterns for all reference states from maximal bias to exponential gradients with mean distances of 30, 90, 180, and 320 $\mu m$, and random. Proliferating tumor cells are red, non-proliferating tumor cells are cyan, and stroma is blue. (b) Peak location, (c) KS statistic comparing proliferating (BrdU+) and non-proliferating (BrdU-) tumor cells, and (d) median of the nearest-stroma distance distribution of proliferating tumor cells for each reference state. Points and boxplots show replicate realizations, and significance annotations are shown relative to the random reference as plotted.

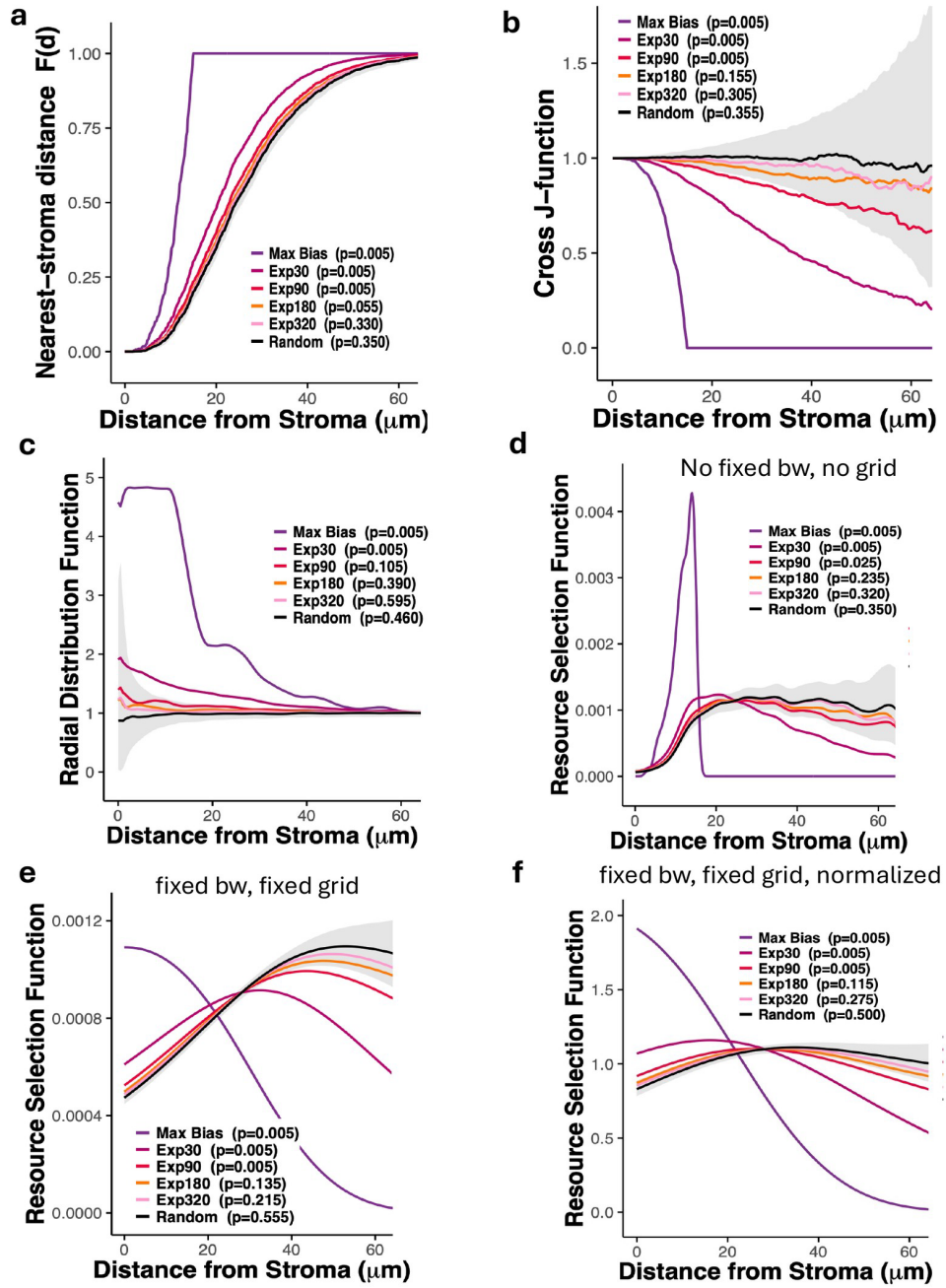

**Supplementary Fig. 3B. High stromal content and high dispersity: global envelopes and RSF estimators.** Global (rank-envelope) versions of the curve statistics for the high-content/high-dispersity specimen of Fig. 4a-e. (a) Cumulative nearest-stroma distance distribution $F(d)$. (b) Cross J-function. (c) Radial distribution function. (d-f) Resource selection function estimated (d) with no fixed bandwidth and no fixed grid, (e) with a fixed bandwidth and a fixed grid, and (f) with a fixed bandwidth and a fixed grid, normalized; in the panel titles, bw denotes the smoothing bandwidth and grid the evaluation grid. Curves show the reference states from maximal bias (Max Bias) to exponential gradients with characteristic distances of 30, 90, 180, and 320 $\mu\text{m}$ (Exp30, Exp90, Exp180, Exp320), and random. The grey band is the global envelope of the random reference; a curve that leaves it at any distance is significant at the global level, and the global $p$-value for each reference state is given in the panel key. Distances extend to about 60 $\mu\text{m}$, the measurable window of this specimen.

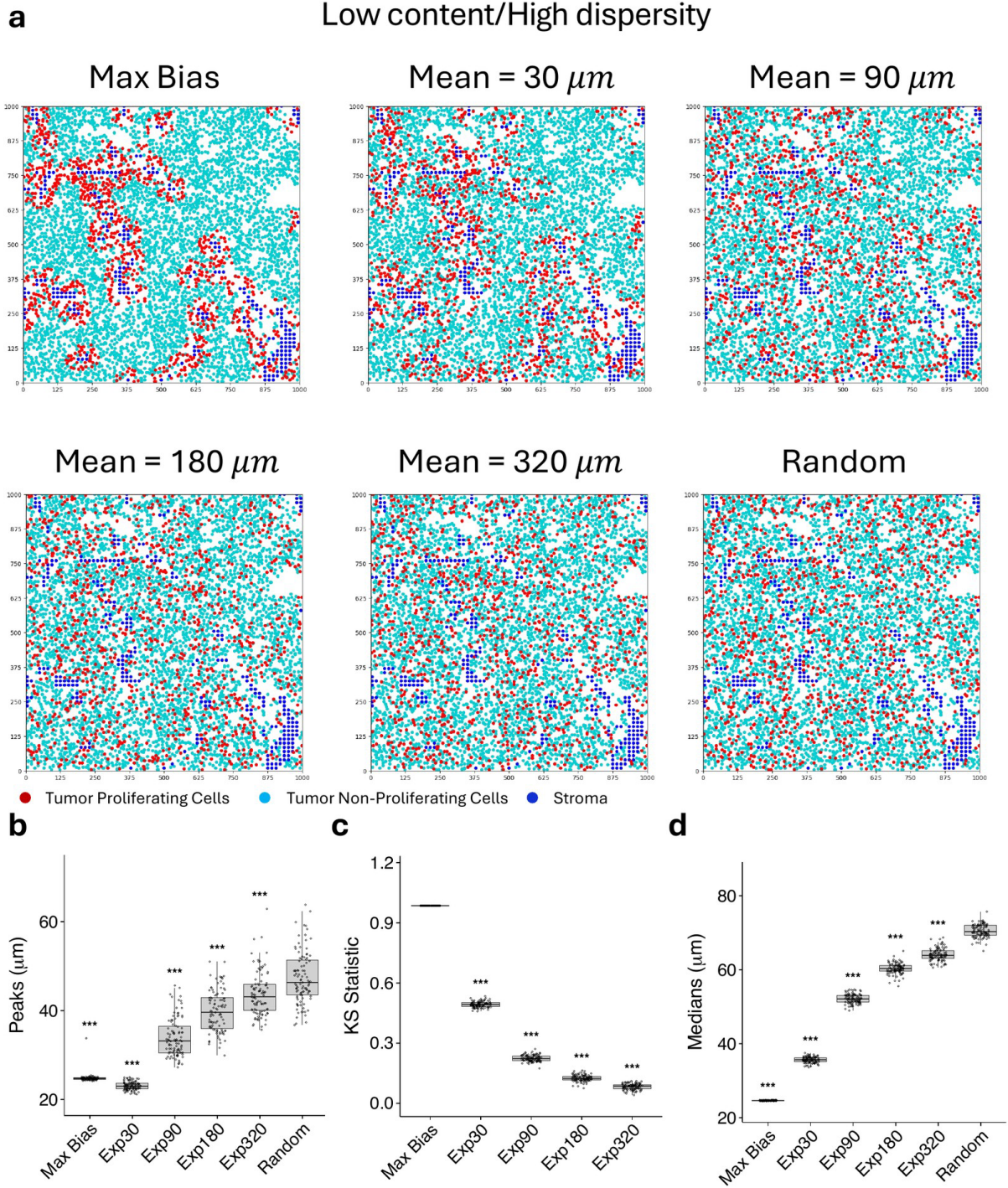

**Supplementary Fig. 4A. Low stromal content and high dispersity: point patterns and distance summaries.** Additional analysis of the low-content/high-dispersity specimen of Fig. 4f-j. (a) Point patterns for all reference states from maximal bias to exponential gradients with mean distances of 30, 90, 180, and 320 $\mu m$, and random. Proliferating tumor cells are red, non-proliferating tumor cells are cyan, and stroma is blue. (b) Peak location, (c) KS statistic comparing proliferating (BrdU+) and non-proliferating (BrdU-) tumor cells, and (d) median of the nearest-stroma distance distribution of proliferating tumor cells for each reference state. Points and boxplots show replicate realizations, and significance annotations are shown relative to the random reference as plotted.

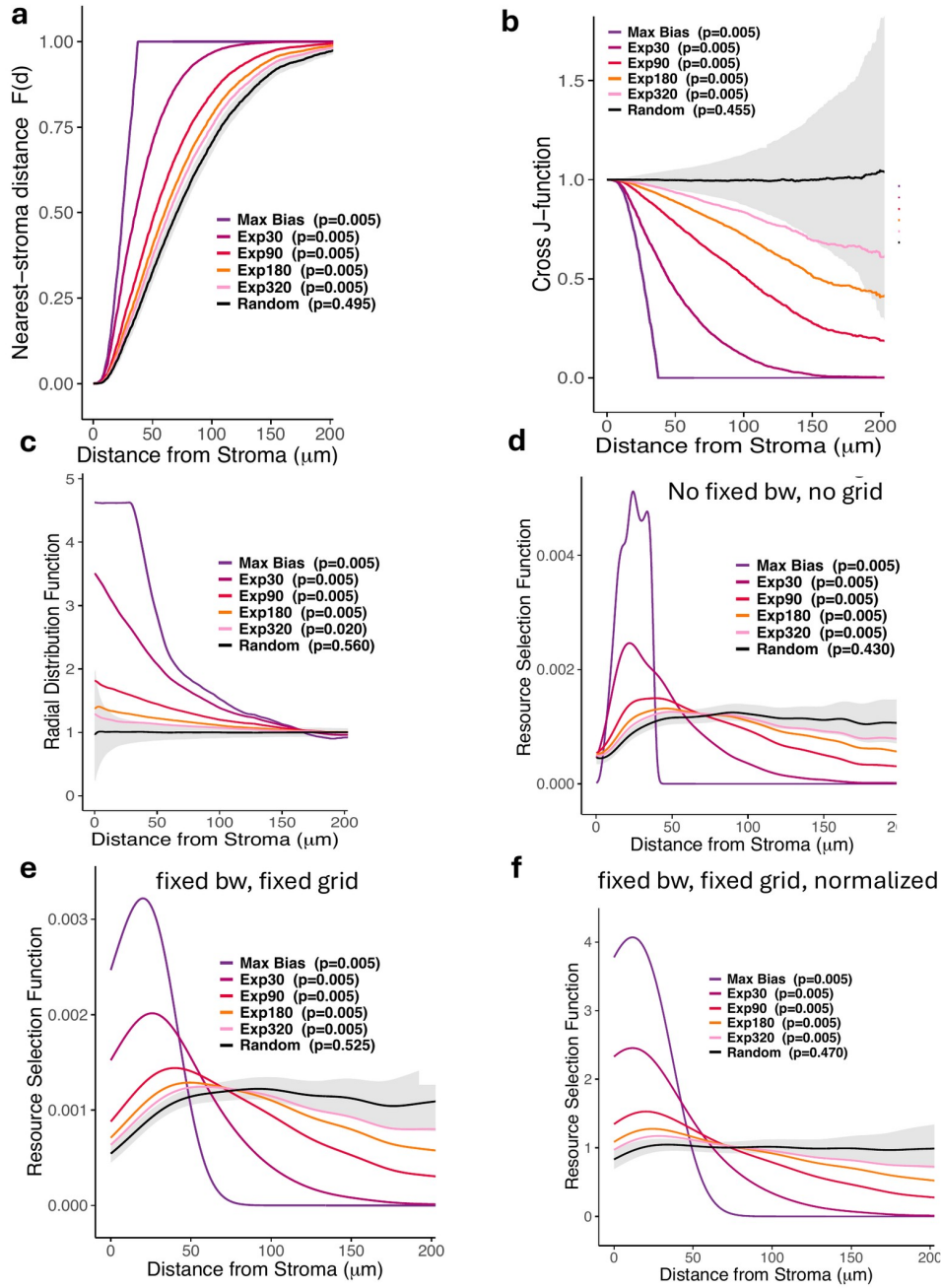

**Supplementary Fig. 4B. Low stromal content and high dispersity: global envelopes and RSF estimators.** Global (rank-envelope) versions of the curve statistics for the low-content/high-dispersity specimen of Fig. 4f-j. (a) Cumulative nearest-stroma distance distribution $F(d)$. (b) Cross J-function. (c) Radial distribution function. (d-f) Resource selection function estimated (d) with no fixed bandwidth and no fixed grid, (e) with a fixed bandwidth and a fixed grid, and (f) with a fixed bandwidth and a fixed grid, normalized; in the panel titles, bw denotes the smoothing bandwidth and grid the evaluation grid. Curves show the reference states from maximal bias (Max Bias) to exponential gradients with characteristic distances of 30, 90, 180, and 320 $\mu\text{m}$ (Exp30, Exp90, Exp180, Exp320), and random. The grey band is the global envelope of the random reference; a curve that leaves it at any distance is significant at the global level, and the global $p$-value for each reference state is given in the panel key. Distances extend to about 200 $\mu\text{m}$, the measurable window of this specimen.

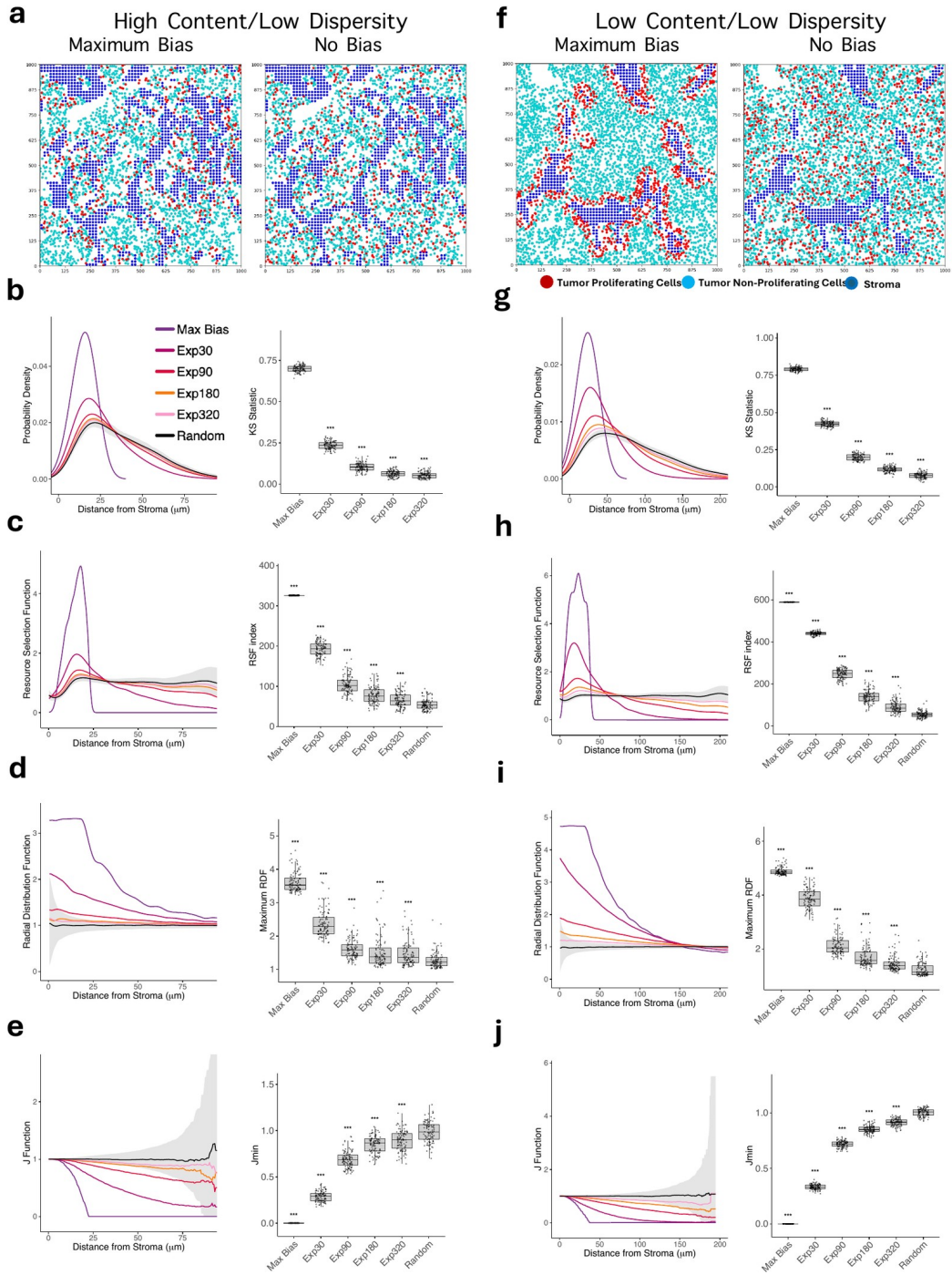

**Supplementary Fig. 5. The two low-dispersy stromal architectures.** High-content/low-dispersy (a-e) and low-content/low-dispersy (f-j) tissues analyzed across the same reference-state ladder, in the layout of Fig. 4. (a,f) Representative point patterns for maximum bias and no bias/random reference. Proliferating tumor cells are red, non-proliferating tumor cells are cyan, and stroma is blue. (b,g) Nearest-stroma distance probability density (left) and KS statistic (right). (c,h) Resource selection function (left) and RSF index (right). (d,i) Radial distribution function (left) and maximum RDF (right). (e,j) Cross J-function (left) and $J_{\min}$ (right). Curves compare maximum bias, Exp30, Exp90, Exp180, Exp320, and random/no bias; grey bands show the random-reference envelope. Points and boxplots show replicate realizations, and significance annotations are shown relative to the random reference as plotted.

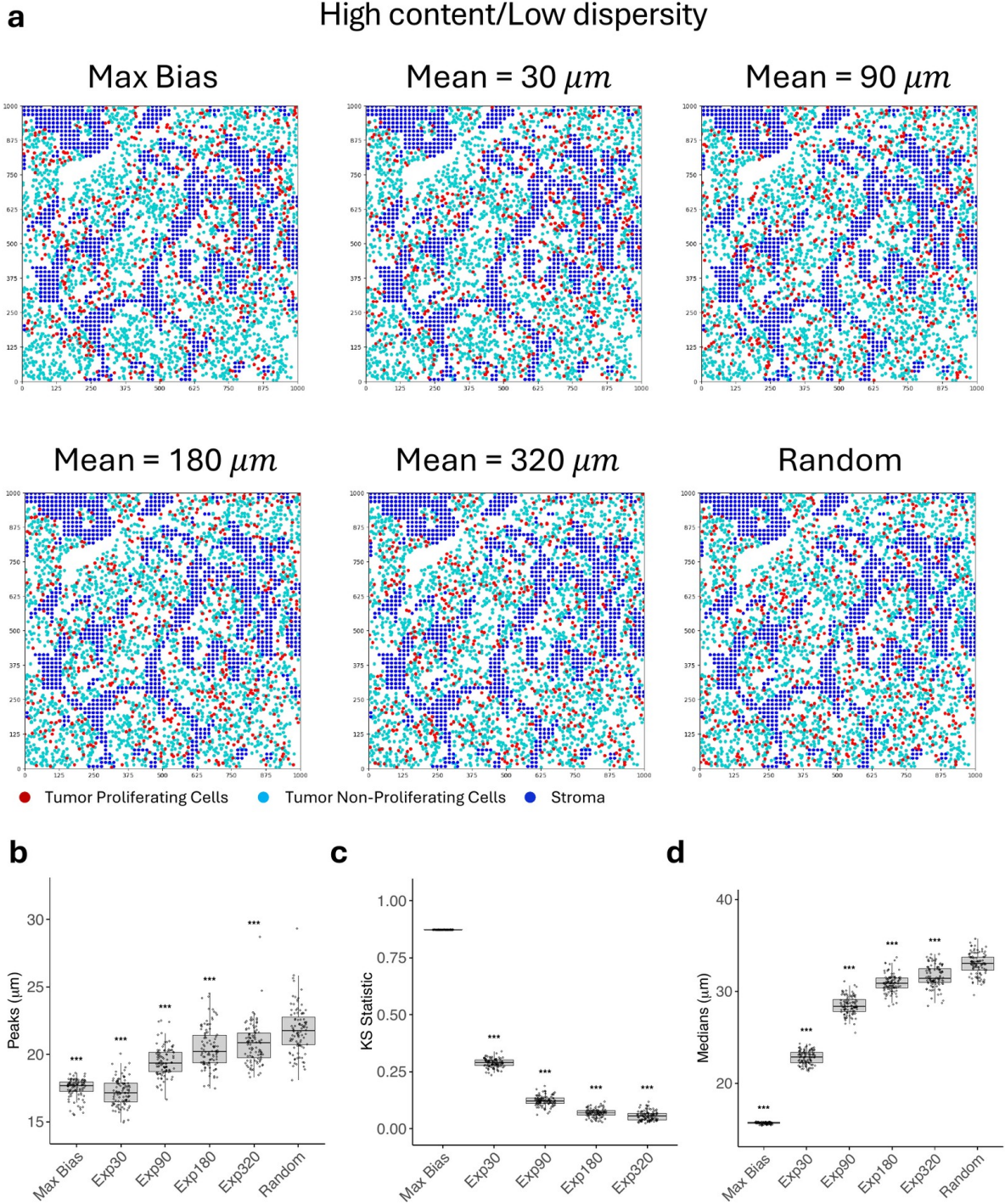

**Supplementary Fig. 6. High stromal content and low dispersity: point patterns and distance summaries.** Additional analysis of the high-content/low-dispersity specimen of Supplementary Fig. 5a-e. (a) Point patterns for all reference states from maximal bias to exponential gradients with mean distances of 30, 90, 180, and 320 $\mu m$, and random. Proliferating tumor cells are red, non-proliferating tumor cells are cyan, and stroma is blue. (b) Peak location, (c) KS statistic comparing proliferating (BrdU+) and non-proliferating (BrdU-) tumor cells, and (d) median of the nearest-stroma distance distribution of proliferating tumor cells for each reference state. Points and boxplots show replicate realizations, and significance annotations are shown relative to the random reference as plotted.

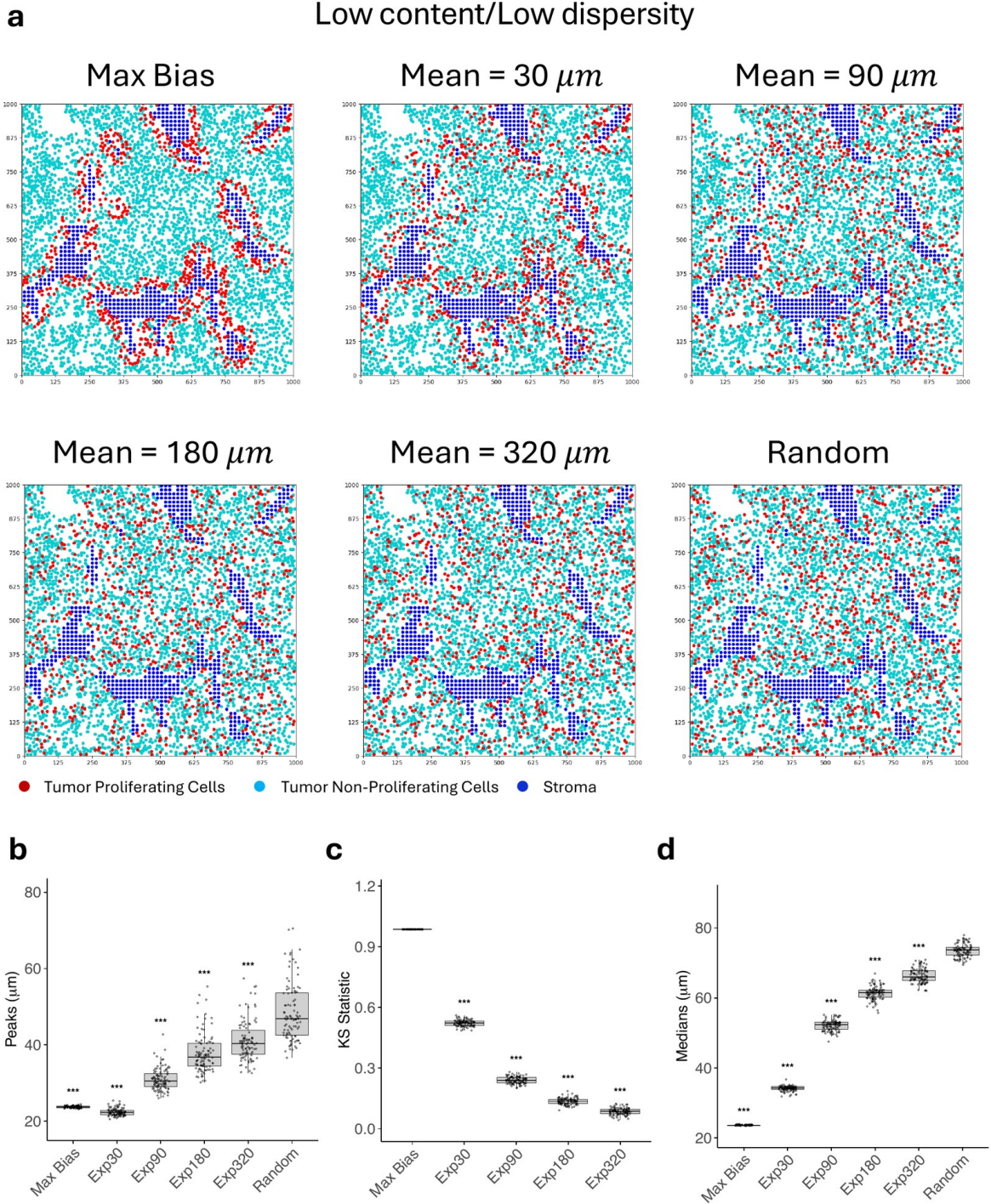

**Supplementary Fig. 7. Low stromal content and low dispersity: point patterns and distance summaries.** Additional analysis of the low-content/low-dispersity specimen of Supplementary Fig. 5f-j. (a) Point patterns for all reference states from maximal bias to exponential gradients with mean distances of 30, 90, 180, and 320 $\mu m$, and random. Proliferating tumor cells are red, non-proliferating tumor cells are cyan, and stroma is blue. (b) Peak location, (c) KS statistic comparing proliferating (BrdU+) and non-proliferating (BrdU-) tumor cells, and (d) median of the nearest-stroma distance distribution of proliferating tumor cells for each reference state. Points and boxplots show replicate realizations, and significance annotations are shown relative to the random reference as plotted.

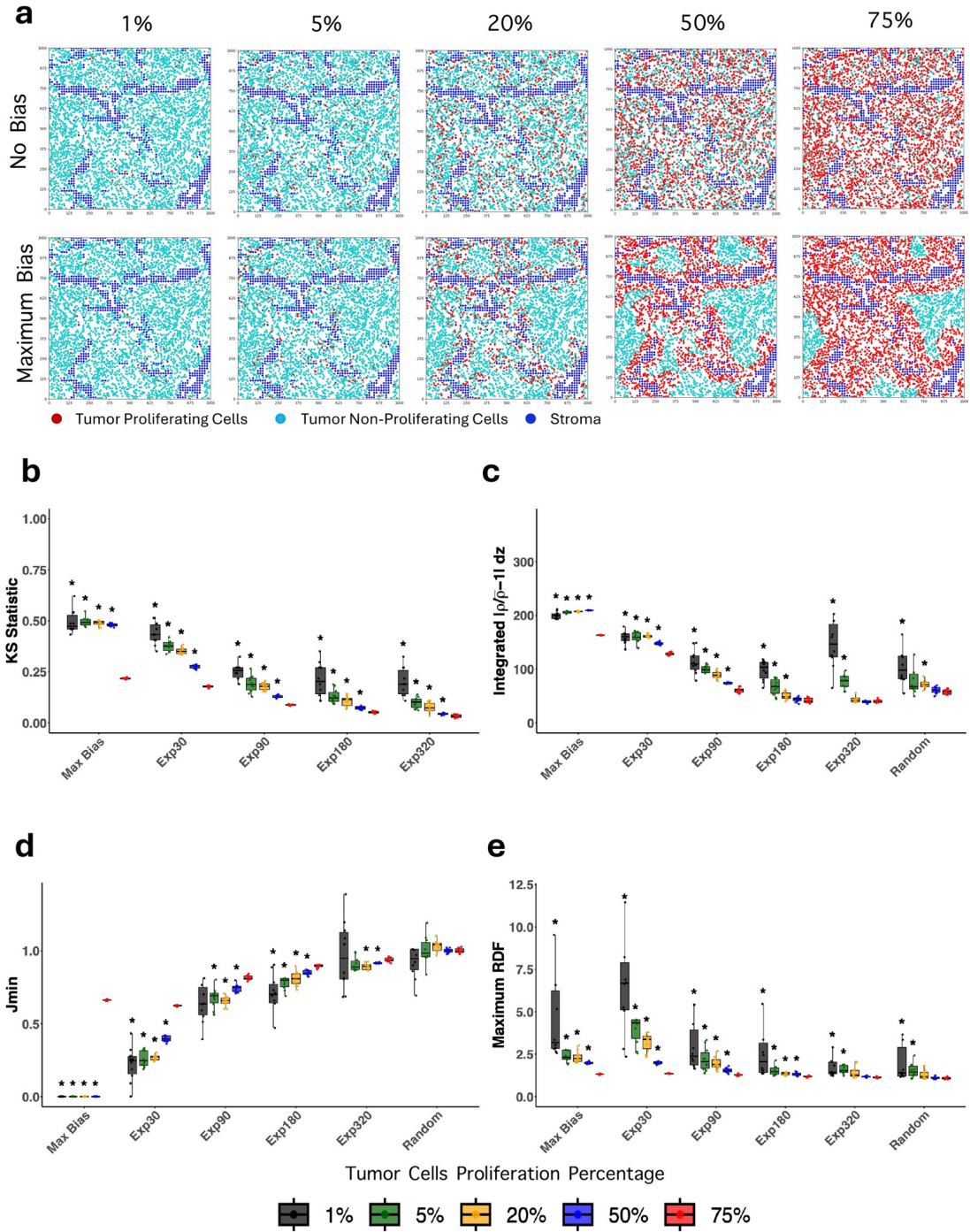

**Supplementary Fig. 8. Proliferative fraction in a first additional stromal architecture.** Proliferative-fraction series as in Fig. 5, for a first additional stromal architecture. (a) Representative point patterns at 1%, 5%, 20%, 50% and 75% proliferating tumor cells under no bias/random assignment (top) and maximum bias (bottom). Proliferating tumor cells are red, non-proliferating tumor cells are cyan, and stroma is blue. (b-e) Summary statistics across the reference-state ladder (maximum bias, Exp30, Exp90, Exp180, Exp320, and random) for each proliferative fraction: (b) KS statistic, (c) RSF index, plotted as the integrated $|\rho/\rho - 1| dz$, (d) $J_{\min}$, and (e) maximum RDF. Colors denote the percentage of proliferating tumor cells as shown in the key.

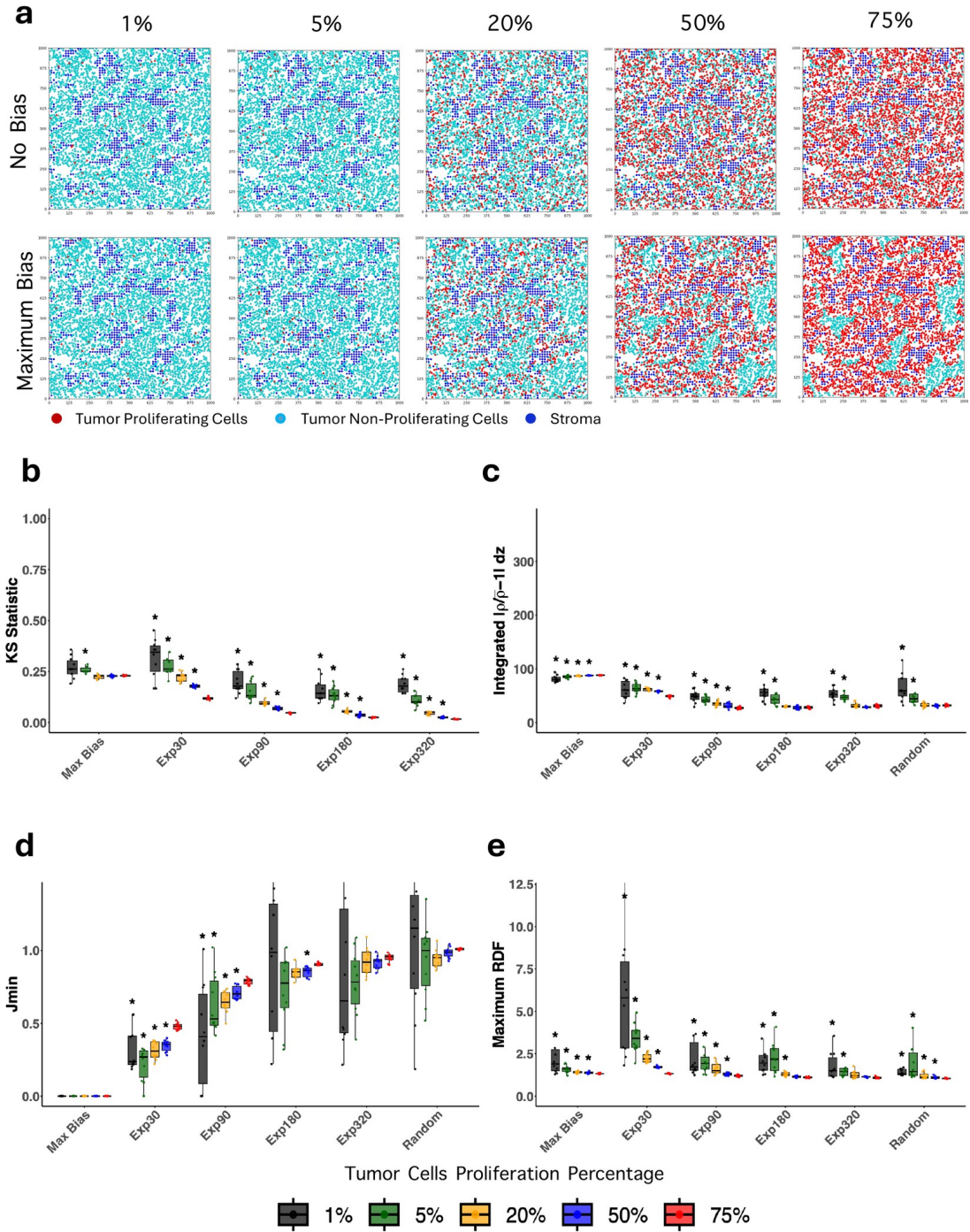

**Supplementary Fig. 9. Proliferative fraction in a second additional stromal architecture.** Proliferative-fraction series as in Fig. 5, for a second additional stromal architecture. (a) Representative point patterns at 1%, 5%, 20%, 50% and 75% proliferating tumor cells under no bias/random assignment (top) and maximum bias (bottom). Proliferating tumor cells are red, non-proliferating tumor cells are cyan, and stroma is blue. (b-e) Summary statistics across the reference-state ladder (maximum bias, Exp30, Exp90, Exp180, Exp320 and random) for each proliferative fraction: (b) KS statistic, (c) RSF index, plotted as the integrated $|\rho/\rho - 1| dz$, (d) $J_{min}$, and (e) maximum RDF. Colors denote the percentage of proliferating tumor cells as shown in the key.

**a**

120 stroma pixels/60
edge pixels (0.5) =
low dispersity

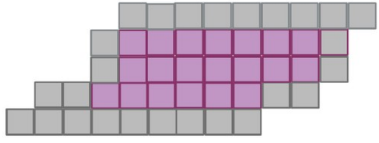

**b**

81 stroma pixels/60
edge pixels (0.74) =
high dispersity

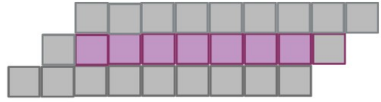

**c**

54 stroma pixels/54
edge pixels (1.0) =
highest dispersity

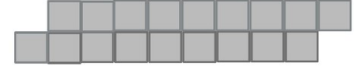

■ Edge stroma pixel

■ Internal stroma pixel

**Supplementary Fig. 10. Stromal edge fraction as a measure of stromal dispersity.** Schematic of the stromal edge-fraction descriptor used to classify stromal architectures (Methods). Edge fraction is the proportion of stromal pixels lying on the stroma-tumor boundary (edge pixels, grey) relative to all stromal pixels; internal stromal pixels that do not contact tumor are purple. (a) A thick, bundled structure of 120 stromal pixels, of which 60 are edge pixels, gives an edge fraction of 0.5 (low dispersity). (b) A thinner structure of 81 stromal pixels with 60 edge pixels gives an edge fraction of 0.74 (high dispersity). (c) A filament of 54 stromal pixels, all of them edge pixels, gives an edge fraction of 1.0 (highest dispersity). Crossed with stromal content, edge fraction defines the four architectural regimes analyzed in Fig. 4 and Supplementary Fig. 5.
